# Flexible neural control of motor units

**DOI:** 10.1101/2021.05.05.442653

**Authors:** Najja J. Marshall, Joshua I. Glaser, Eric M. Trautmann, Elom A. Amematsro, Sean M. Perkins, Michael N. Shadlen, L.F. Abbott, John P. Cunningham, Mark M. Churchland

## Abstract

Voluntary movement requires communication from cortex to the spinal cord, where a dedicated pool of motor units (MUs) activates each muscle. The canonical description of MU function rests upon two foundational tenets. First, cortex cannot control MUs independently but supplies each pool with a common drive. Second, MUs are recruited in a rigid fashion that largely accords with Henneman’s size principle. While this paradigm has considerable empirical support, a direct test requires simultaneous observations of many MUs across diverse force profiles. We developed an isometric task that allowed stable MU recordings even during rapidly changing forces. MU activity patterns were surprisingly behavior-dependent. MU activity could not be accurately described as reflecting common drive, but could be captured by assuming multiple drives. Neuropixels probe recordings revealed that, consistent with the requirements of flexible control, the motor cortex population response displays a great many degrees of freedom. Neighboring cortical sites recruited different MUs. Thus, MU activity is flexibly controlled to meet task demands, and cortex may contribute to this ability.

## Introduction

Primates produce myriad behaviors, from acrobatic maneuvers to object manipulation, all requiring precise neural control of muscles. A single muscle is controlled by a motor neuron pool (MNP) containing hundreds of α-motoneurons, each innervating a unique subset of muscle fibers^1^. A motoneuron and the fibers it innervates constitute a “motor unit” (MU), the smallest controllable element of the neuromuscular system^2^. MUs are highly heterogeneous^3^, differing in size (large MUs innervate more fibers)^4,5^, duration of generated force^6^, and muscle length where force becomes maximal^7^. Nearly every functional property of muscle fibers varies, across MUs, over a 2-10 fold range^3,6,8^.

Optimality suggests flexibly using MUs best suited to each situation^9–11^. Yet flexibility necessitates intelligently controlling many degrees of freedom. A simpler alternative is a rigid strategy that can approximate optimality. Supported by nearly a century of research, rigid recruitment is the canonical conception of MU control^12–14^. With limited exceptions, MUs are believed to be recruited in a consistent order^15,16^ from smallest to largest according to Henneman’s size principle^17–23^. Fixed recruitment is attributed to intrinsic motoneuron properties^2,13,14^. Small-to-large recruitment minimizes force fluctuations^24^ and energy expenditure^25^. It is also computationally efficient^26^: one ‘common drive’ force command determines every MU’s activity^27,28^. This accords with the broader belief that control is limited to relatively few ‘synergies’ (e.g., a ‘flexion-synergy’ that activates multiple flexor muscles)^29–34^. Henneman found small-to-large recruitment regardless of whether input was delivered via electrical stimulation of dorsal roots^17^; electrical stimulation of the motor cortex, basal ganglia, cerebellum, or brain stem^35^; or mechanical activation of reflex circuits^18–20^.

Rigid control is thought to simplify control (**Fig. 1a**). If “the brain cannot selectively activate specific motor units^13^”, it is freed from the need to do so. The activity of each MU is a fixed function of a unified force command – typically one command per muscle, though a muscle with more than one mechanical action may be driven by more than one synergy. In principle, rigid control could use any set of fixed ‘link’ functions. In practice, link functions are thought to yield small-to-large recruitment, with limited well-established exceptions. The most trivial exception is that apparent changes in recruitment order occur during ballistic contractions, due to fixed conduction-velocity differences^12^. Each MU’s activity is still proposed to be a fixed function of force, just with slightly different latencies^36,37^. A second exception is that, for muscles with multiple mechanical actions, the size principle holds only within each action^11,38,39^. Elbow flexion, forearm supination, and humerus exorotation each activate a somewhat different region of the biceps^40^, and each displays its own size-based recruitment^41,42^. It is hypothesized that each mechanical action has its own common drive, which overlap imperfectly within the MNP^41^ (**Fig. 1a**, *middle*) and are preferentially reflected by different nerve branches^43^. Recruitment is still considered rigid^44^ because it remains a fixed function of simple commands (one per mechanical action). Another well-established phenomenon is that cell-intrinsic mechanisms cause some MUs to rotate on and off (on a timescale of many seconds) to combat fatigue^45^. Thus, although the size principle is obeyed on average, it may not perfectly describe the response on every trial. It is cautioned that these phenomena can be misinterpreted as flexibility^36,44,46,47^ and must be controlled for.

**Figure 1.**
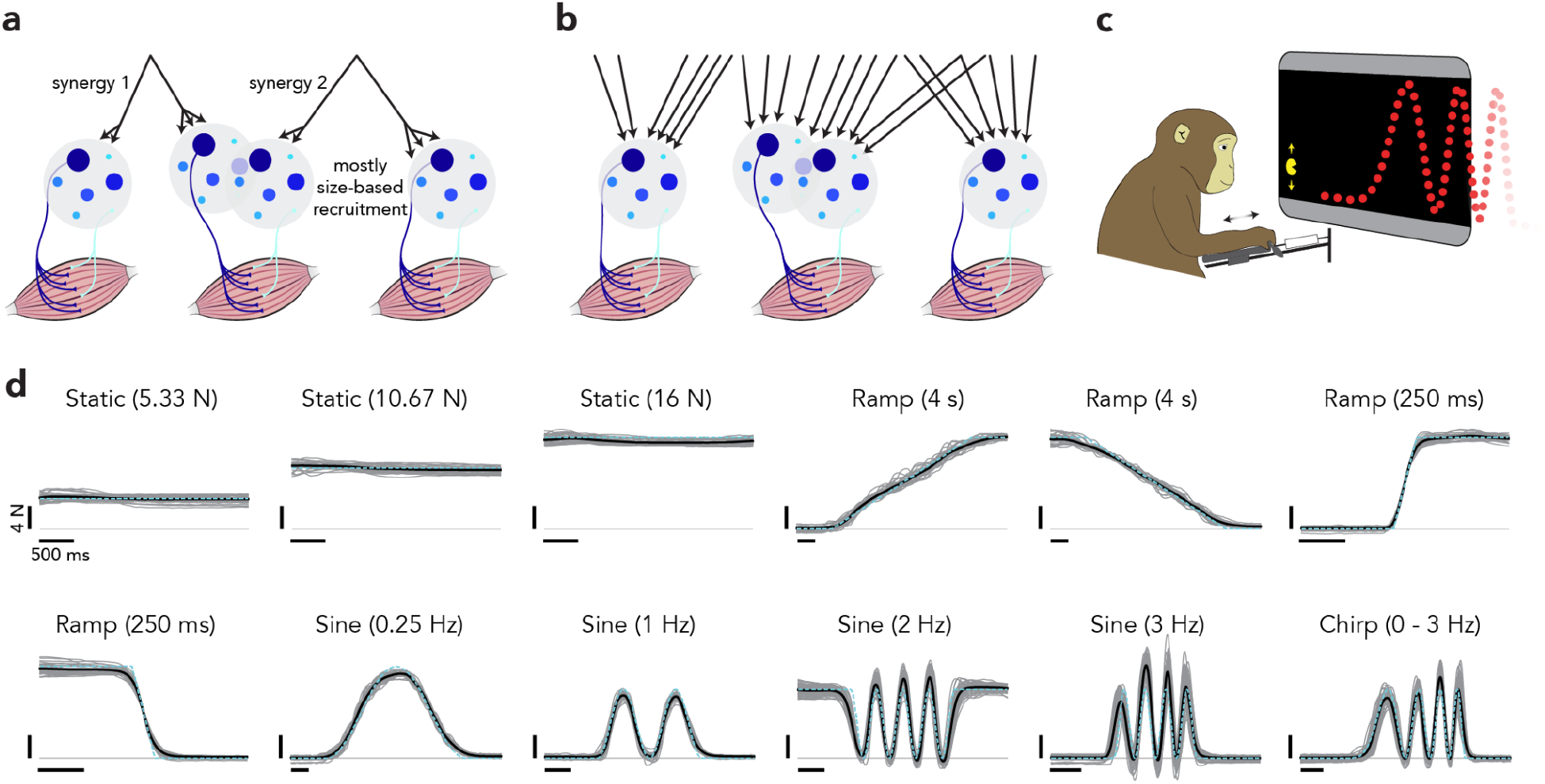
Rigid versus flexible motor unit control and experimental setup. (**a**) Rigid control schematic. The activity of many MUs is determined by a small number of force commands, perhaps ‘synergies’ that control particular mechanical actions (i.e., an elbow flexion synergy and a forearm supination synergy). Thus, the number of controlled degrees of freedom is less than or equal to the number of motor neuron pools. Each MU’s activity is a function of that small number of degrees of freedom, and those functions typically enforce size-based recruitment. (**b**) Flexible control schematic. Each motor neuron pool is controlled by multiple degrees of freedom, such that recruitment can be size-based but can also be flexibly altered when other solutions are preferable. (**c**) Task schematic. A monkey modulated the force generated against a load cell to control Pac-Man’s vertical position and intercept a scrolling dot path. (**d**) Single-trial (*gray*), trial-averaged (*black*), and target (*cyan*) forces from one session. Vertical scale: 4 N. Horizontal scale: 500 ms.

The opposite of rigid control -- highly flexible control that tailors recruitment by leveraging many degrees of freedom (**Fig. 1b**) -- has rarely been considered a viable possibility. Yet indirect measures such as EMG frequency content suggest that recruitment changes within the gait cycle in rats^48,49^ and across cycling speeds in humans^50^, suggesting more flexibility than generally appreciated^51^. Still, most studies argue that speed-based recruitment is empirically absent^37,52,53^ and/or of limited benefit^54^. The consensus has thus been that recruitment is independent of speed^13,44,55,56^. A possible exception is fast lengthening contractions^10^, but studies disagree^57^ and this appears not to be a general property^55^. Another possibility is that latent flexibility may be revealed via biofeedback training^46,47,58–61^. However, it remains unclear whether subjects truly display flexible control or simply leverage known size-principle exceptions; biofeedback-based control appears quite limited^62^. Thomas^46^ found biofeedback-based flexibility only in muscles with multiple actions and argued it is a trivial consequence of subjects learning to move differently (e.g., supinating versus flexing). The prevailing view is thus that flexibility is scant^44,55,56^ or non-existent^44^. Even those most positive regarding potential flexibility suggest that orderly recruitment seems “broadly adhered to in the majority of cases^54^”. Yet many have stressed the need for further examination using improved recording techniques^51^ and a broader range of behavioral conditions^44^.

Because rigid and flexible recruitment make divergent predictions regarding what cortex can control, we first examined recruitment following cortical stimulation. Then, to ascertain which behavioral situations might reveal flexibility, we used optimization to discern when rigid and flexible control make divergent predictions. We employed a highly practiced force-tracking task with a broad range of behavioral conditions, including those predicted to be diagnostic. Because the challenge of simultaneously recording many MUs^63^ has limited studies of swiftly changing forces^48–50^, we used modified methods for recording and sorting spikes from many neighboring MUs. Additionally, we used Neuropixels recordings to assess the degrees of freedom naturally present in cortical population activity.

## Results

### Pac-Man Task and EMG recordings

We trained a rhesus macaque to perform an isometric force-tracking task. The monkey modulated force to control a ‘Pac-Man’ icon and intercept scrolling dots (**Fig. 1c**). We instructed force profiles (**Fig. 1d**) via the dot path. Following extensive training, empirical forces (single trials in *gray*) closely matched target force profiles (*cyan*). We conducted three experiment types, each using dedicated sessions (38 ~1 hour sessions performed on separate days). Microstimulation experiments perturbed descending commands. Dynamic experiments employed diverse force profiles. Muscle-length experiments investigated MU recruitment across muscle lengths.

In each session, we recorded from multiple custom-modified percutaneous thin-wire electrodes closely clustered within the head of one muscle. EMG spike-sorting is notoriously difficult during dynamic tasks; movement threatens recording stability and vigorous activity produces superimposed action-potential waveforms^64^. Three factors enabled identification of spikes from many individual MUs: the isometric task facilitated stable recordings, intensity could be titrated via task gain, and each MU produced a unique complex waveform typically spanning many channels (**Fig. S1**). Spike waveforms were identified by adapting recent spike-sorting^65–67^ advances, including methods for resolving superimposed waveforms^66^ (Supp. Materials).

We isolated 3-21 MUs per session (356 total MUs). To eliminate any possibility that recruitment could mistakenly appear flexible due to known properties of multifunctional muscles, forces were always generated with the same mechanical action. Analyses considered only simultaneously recorded neighboring MUs (we did not compare MUs recorded in different parts of the muscle) and recordings included non-multifunctional muscles. This is important given the long-standing caution that known properties of rigid recruitment should not be misinterpreted as flexibility^36,44,56^.

### Cortical perturbations

Rigid MU control is thought to arise from a spinally enforced recruitment mechanism^19^. Artificially perturbing cortex should therefore modulate overall MNP activity while preserving orderly recruitment^35^. In contrast, fully optimal control would likely require participation from areas aware of the overall action and context. If so, cortex might have the capacity to alter recruitment. It is argued that this does not occur^35^. However, the possibility is suggested by the finding that pyramidal tract stimulation produces different synaptic inputs to small and large MUs in anesthetized cats^68^ and by mapping experiments in baboons^69^.

We re-examined this topic by delivering microstimulation (57 ms, 333 Hz) through a linear array (100 um interelectrode distance) in sulcal motor cortex (M1) (**Fig. 2a**). Recordings were made from the *deltoid*, *triceps*, and *pectoralis* over 18 daily sessions (average of 370 trials per session, 3-21 simultaneously recorded MUs, 183 total MUs). We optimized probe location to activate the recorded muscle, and chose a subset of neighboring electrodes for stimulation. Stimulation often activated only part of a muscle, and recordings were targeted to that subregion. Stimulation was delivered during static forces. A slowly increasing force ramp was performed without stimulation.

**Figure 2.**
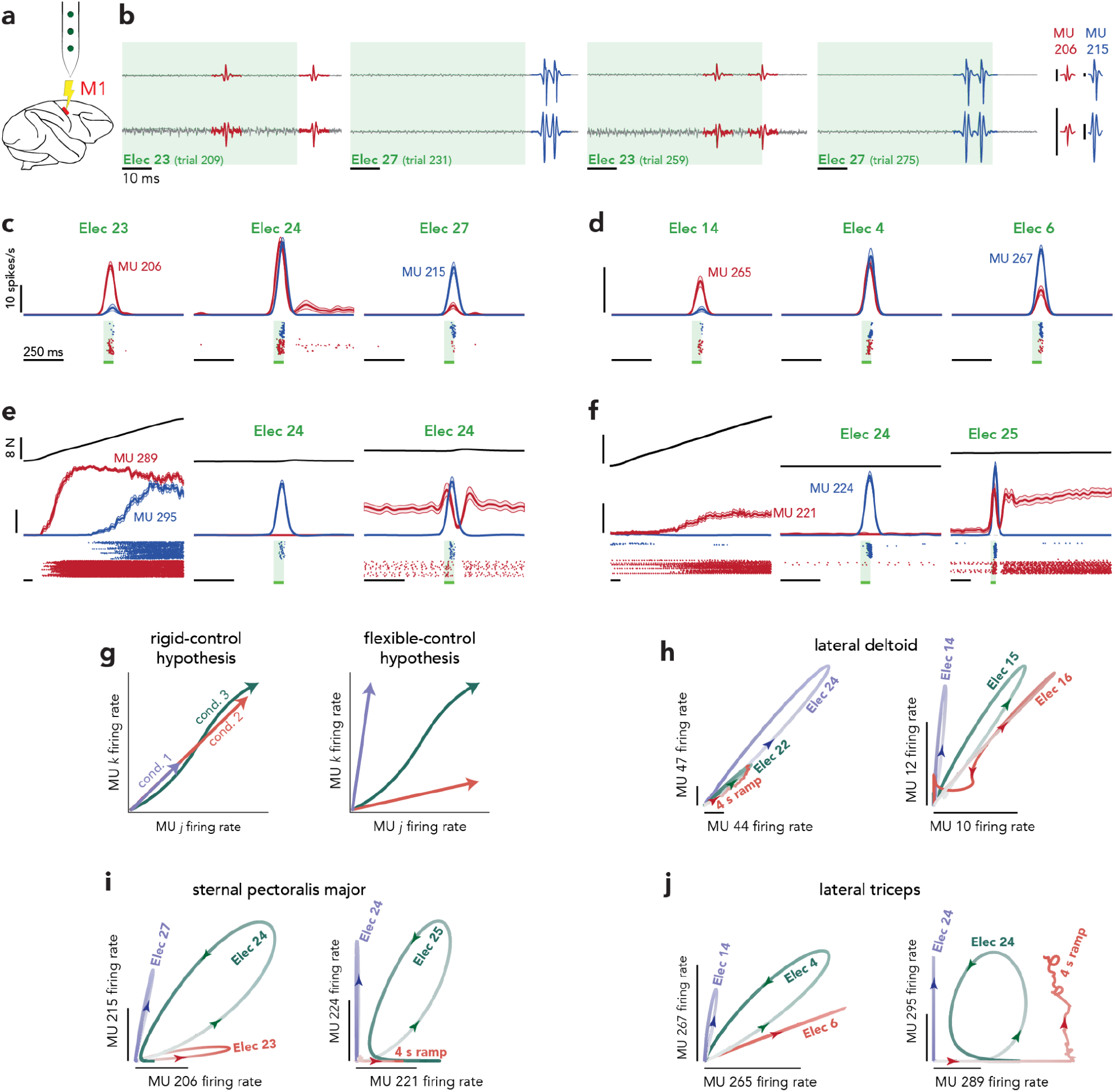
Stimulation experiments. (**a**) Microstimulation was delivered through a selection of electrodes on a linear array. (**b**) EMG voltages for four example trials and 2 (of 8) EMG recording channels. To aid visualization of MU waveforms with different magnitudes, vertical scales are customized to the stimulated electrode. Stimulation was delivered during the *green-shaded* region, during which a small stimulation artifact can be seen. *Right*. spike templates for the two MUs present in the example trials. Vertical scale: 50 standard deviations of the background noise (ignoring any stimulation artifact). Expanded vertical scale for MU206 to match that in example trials. (**c**) Example responses of two MUs during stimulation on three electrodes. Traces plot firing rates (mean and SEM). Rasters show MU spikes, separately for the two MUs, on every trial. (**d**) As in **c** but during a different session. (**e**) Example MU responses during the slow ramp and stimulation during two different baseline force levels. Format as above, with mean force plotted at top. (**f**) Similar format for a different session. Responses are shown during the slow ramp and stimulation on two sites. (**g**) State space predictions for rigid and flexible MU control. The firing rate of one MU plotted against that of another should fall along the same 1-dimensional monotonic manifold for all conditions if control is rigid (*left*) or along different manifolds if control is flexible (*right*). (**h**) State space plots, each showing two MUs recorded from the lateral deltoid during stimulation on three electrodes. (***i***) Same for sternal pectoralis major. Responses are from three stimulation sites (*left*) or two sites and during the slow force ramp (*right). (**j***), Same for lateral triceps.

Under rigid control, the MU population response should be compatible with common drive and MU-specific link functions. In testing this hypothesis, a challenge is that common drive and the link functions are unknown hypothesized quantities. One can thus reject the hypothesis of rigid control only if responses cannot be explained by any plausible common drive and link functions. We pursue this goal directly in a later section. To build towards that goal, we first consider MU pairs. Purely pairwise comparisons are conservative; many pairwise responses that suggest flexible control are also explainable under rigid control if one is willing to make generous assumptions regarding the link functions. Nevertheless, if control is truly flexible, unmistakable departures from rigid control should sometimes be observable at the level of MU pairs. One such departure is illustrated in **Fig. 2b** at the level of raw voltage traces (only two recording channels shown for simplicity). Stimulation recruited two neighboring *pectoralis* MUs, with waveforms distinguishable based on shape and amplitude (MU206 is much smaller amplitude and shown on an expanded scale). Electrode 23 recruits MU206 and electrode 27 recruits MU215. Across trials, it was consistently true that electrode 23 primarily recruited MU206 and electrode 27 primarily recruited MU215 (**Fig. 2c**). Electrode 24 (physically between 23 and 27) recruited both MU206 and MU215.

The activity in **Fig. 2b,c** is incompatible with electrodes 23 and 27 recruiting the same common drive. A similar effect is shown for two *triceps* MUs in **Fig. 2d**. In 13/18 sessions, the strength of activation across MUs depended on which electrode delivered the stimulation (two-way ANOVA; p < 0.001, adjusted for multiple comparisons, for an interaction between MU identity and stimulating electrode; n = 11-77 trials per condition; session average = 29). While suggestive, we stress that an ANOVA identifies one specific effect type that is likely (but not necessarily) a deviation from rigid control. A more focused and stringent test is presented in the next section.

Cortical stimulation also yielded responses that departed from the putatively “standard” recruitment during a slowly increasing force ramp. During the ramp (**Fig. 2e**, *left*), the joint activity of MU289 (*red*) and MU295 (*blue*) are explainable by link functions that become positive at low (MU289) and medium (MU295) forces. Yet electrode 24 activated MU295, but not MU289 (**Fig. 2e**, *middle*). When MU289 was already active during static force production (**Fig. 2e**, *right*), stimulation had an effect consistent neither with common drive (it differed for the two MUs) nor with recruitment during the ramp (where MU289 was always more active).

Examples illustrate effect range and robustness (SEMs are small) and allow one to consider whether there might exist alternative explanations. It can be useful to do so at the level of rates and single-trial spike trains (rasters at bottom) before subsequent quantification. One potential trivial source of apparent departures from rigid control is small latency differences amongst MUs^36^. Inspection reveals this cannot explain most stimulation-induced departures. For example, in the left two panels of **Fig. 2e** and **2f**, activity departs from rigid control not because of a latency difference between MUs, but because the force ramp mostly recruits one MU while stimulation recruits another. At the same time, small latency differences definitely exist: e.g., MU206 rises ~20 ms earlier than MU215 (**Fig. 2c**). Subsequent quantification is thus careful to take this into account.

We saw little evidence of general fatigue over the course of a session (**Fig. S2b**), presumably because force levels were modest. Occasionally, the classic fatigue-combatting rotations^70^ produced ‘streaky’ spike rasters (**Fig. 2f**, *red*) but this was not the cause of departures from rigid control seen in trial-averaged rates. To minimize the impact of both general fatigue and fatigue-combatting rotations on trial-averaged responses, trials for all conditions (including stimulation sites) were interleaved. For example, consider the red rasters in **Fig. 2e**. The trials where the ramp produced a robust response were interleaved with trials where electrode 24 produced no response. Occasionally, cortical perturbations produced hysteresis (**Fig. 2f**, *right*), likely reflecting persistent inward currents^71^. Unlike the immediate effect of stimulation, hysteresis rarely altered recruitment order relative to natural behavior, consistent with the finding that persistent inward currents rarely alter recruitment order because they are most prevalent in small MUs^56^.

### State-space predictions of rigid control

The hypothesis of rigid control is particularly easy to express in a state space where each MU’s activity contributes an axis. Under rigid control, all MUs have activity that is a monotonic function of common drive^72–74^. Thus, with increasing force, activity should move farther from the origin, tracing a monotonic one-dimensional (1D) manifold (**Fig. 2g**, *left*). The manifold may curve (due to nonlinear link functions) but greater drive should cause all rates to increase (or remain unchanged if maximal or not yet recruited). In contrast, flexible control predicts recruitment patterns unconfined to a monotonic 1D manifold (**Fig. 2g**, *right*) and that can ‘fill up’ much more of the state space. To visualize this, we first consider the joint activity of MU pairs. Responses sometimes adhered to classical predictions (**Fig. 2h**, *left*). Yet stimulation could drive deviations from a monotonic 1D manifold (**Fig. 2h**, *right;* **Fig. 2i, j**), both when comparing among electrodes and when comparing with the ramp. Brief ‘loops’ are compatible with rigid control because they likely reflect small latency differences amongst MUs^36^. In contrast, entirely different trajectory directions across conditions indicate departures from rigid control.

Standard statistical tests (e.g., the ANOVA above) can reveal when responses are statistically different among MUs. However, the key question is not whether responses differ, but whether differences can be explained by different link functions operating upon the same common drive. In a subsequent section we address this question by fitting a model. While conceptually simple, model fitting requires optimization of many free parameters. Thus, we first use an alternative strategy: asking whether there exist moments during population-state trajectories that are logically inconsistent with rigid control, regardless of choice of link functions / common drive.

We first applied an ‘MU displacement’ metric (**Fig. 3a**). Suppose two MUs have rates of 10 spikes/s at time *t*. If at time *t′* (possibly within a different condition) both have higher rates (*left*), one can connect them with a monotonic manifold (*dashed line*). If rates change in opposition (one increasing, one decreasing, *right*), they can no longer be connected by a monotonic 1D manifold. ‘Displacement’ is the smallest alteration (*blue*), in either MU’s rate, that allows states to be connected by monotonic manifold. For each empirical population response, we identified the MU pair causing the largest displacement, then minimized this displacement by allowing temporal shifts to eliminate latency-based effects. Any remaining displacement was the maximal displacement for that session. For 16/18 sessions, maximum displacement was higher than expected from sampling error (**Fig. 3b**, p<0.001, two-sample t-test). Displacements were larger when considering population trajectories across all stimulation sites (“all-stim”) versus each site alone (“1-stim”). This agrees with what can be seen in the examples: large deviations from rigid control are clearest when comparing across stimulation sites. However, displacement was also sometimes non-negligible even within a single site, which occurs if stimulation drives two MUs with opposing polarities or different temporal patterns (e.g., **Fig. 2e**, *right*).

**Figure 3.**
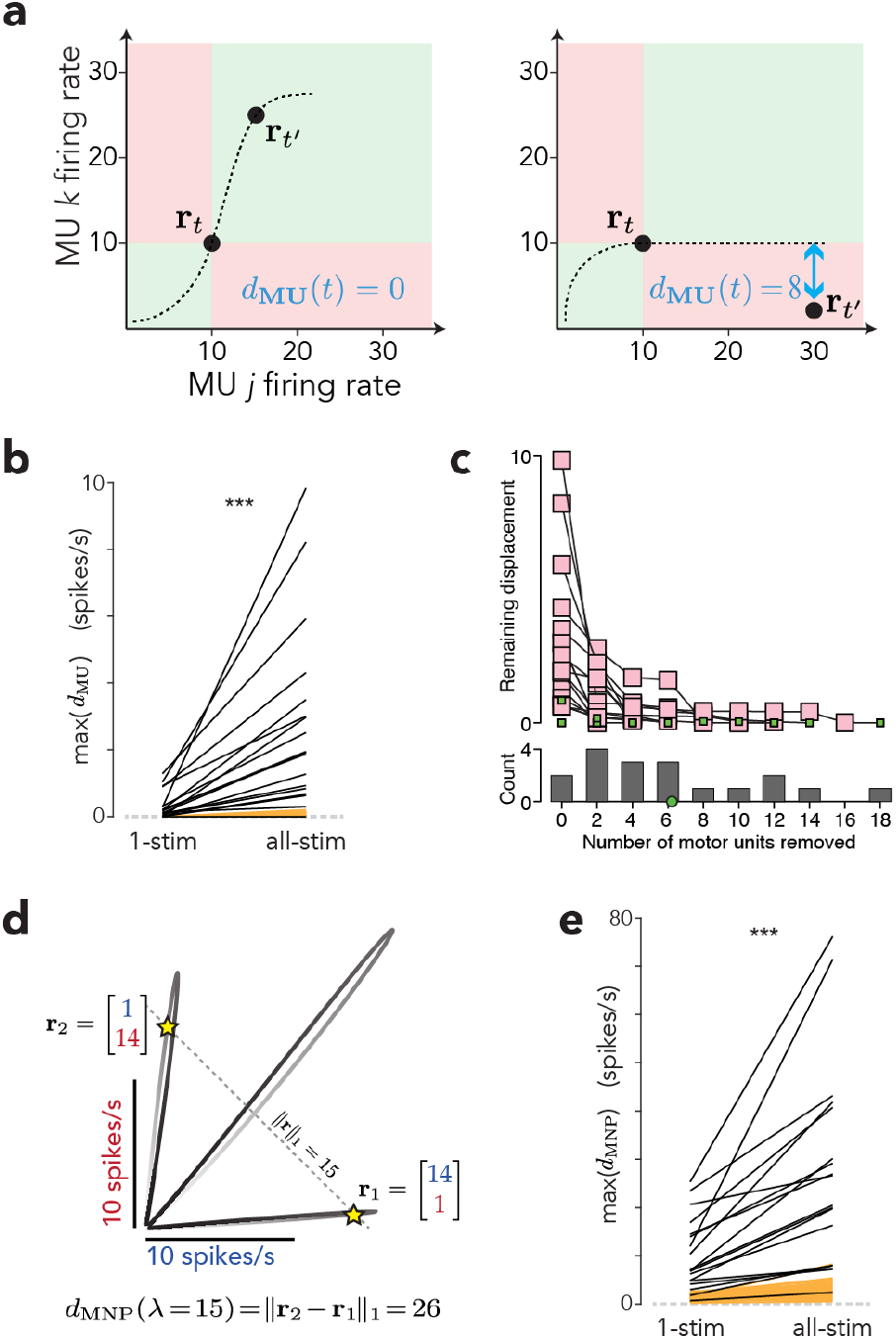
Quantifications of flexible control. (**a**) Schematic illustrating MU displacement (*d*_MU_) for a two-dimensional population state and two times. *Left*. a monotonic manifold can pass through **r***_t_* and **r***_t′_*. Thus, *d*_MU_(*t*) = 0. *Right*. Any monotonic manifold passing through **r***_t_* is restricted to the green zone, and thus cannot come closer than 8 spike/s to **r***_t′_*. Thus, *d*_MU_(*t*) = 8. (**b**) Maximum displacement (one black line per session). For ‘1-stim’, displacement was computed considering only times within the response for each stimulation site. The maximum was then taken across sites. For ‘all-stim’, the maximum displacement was computed across all times and stimulation sites. ‘All-stim’ displacement was significantly larger (two-sample t-test, p < 0.001, N=18 sessions). Both were significantly larger than the displacement expected due to sampling error alone (*orange*, ten resampled artificial populations per empirical dataset). (**c**) Maximum displacement (as for ‘all-stim’) after repeatedly removing the MU pair that caused the largest displacement. Each trace corresponds to one session, and ends when remaining displacement is no larger than the largest across the ten resampled populations. Gray histogram plots the distribution how many MUs had to be removed to reach that point (mean in *green*). (**d**) Illustration of MNP dispersion metric for two MUs. The three trajectories were driven by stimulation at three cortical sites. The line defined by ║r║_1_ = 15 intercepts these trajectories at multiple states, with **r**_1_ and **r**_2_ being most distant. The MNP dispersion for λ = 15 is the L1-norm of the difference between **r**_1_ and **r**_2_ (**e**) Maximum (across λ) dispersion for each session. ‘All-stim’ dispersion was significantly larger (two-sample t-test, p < 0.001, N=18 sessions). Both were significantly larger than the dispersion expected due to sampling error alone (*orange*, ten resampled artificial populations per empirical dataset). Results are shown for conditions where stimulation was delivered as the monkey held a low static force, and were nearly identical for a higher static force.

We extended the above analysis by removing, from the population, the MU pair responsible for the largest displacement. We recomputed displacement, then repeated the process until remaining displacement was no larger than the largest expected from sampling error alone (**Fig. 3c**). On average it was necessary to remove 6.2 MUs, out of an average of 10.2 per session.

The second metric, ‘MNP dispersion’, leverages the fact that common drive putatively determines the population state and thus the L1-norm, ║r║_1_. A given value of ║r║_1_ corresponds to a given level of common drive and thus a unique population state. Geometrically, a monotonic manifold should not cross the diagonal defined by a given ║r║_1_ in multiple places (**Fig. 3d**). We thus define MNP dispersion, *d*_MNP_(λ) as the maximum distance between all population states with norm λ. For each session we computed the maximum dispersion (across λ), then minimized that value by allowing latency shifts. Dispersion was higher than expected given sampling error for all but one session (**Fig. 3e**) and was larger when considering all stimulation sites than when considering each individually.

### Optimal motor unit recruitment

Is the capacity for flexible MU recruitment used during behavior? Because larger MUs tend to generate shorter-lived forces^75^, a venerable suggestion is that, if flexibility exists, it might reveal itself when comparing slow versus fast movements^10,48–50^. The prevailing view is that this does not occur^76^. Recruitment appears mostly preserved during slowly and quickly increasing force ramps^36,37^, with the exception of trivial effects due to latency. Yet some studies have found evidence for flexible recruitment^10,50^ using force profiles other than steps. We reasoned that any revisitation of this question should be informed by expectations regarding which force profiles are mostly likely to evoke altered recruitment.

We developed a simplified computational model, inspired by prior work that established size-based recruitment as the optimal fixed strategy^24–26^. We ask what would occur if recruitment were infinitely flexible and optimized for each force profile. We modeled the force produced by an idealized MNP with *N* MUs as

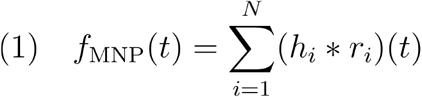

where *h_i_*(*t*) and *r_i_*(*t*) are the twitch response and trial-averaged firing rate of the *i*^th^ MU. We used the model of Fuglevand *et al*. to simulate twitch responses^73^, which were bigger and briefer for larger MUs. For each target force profile, we derived the set of MU rates that minimized mean-squared error (both average error and trial-to-trial variability) between *f*_MNP_(*t*) and intended force (**Fig. 4a**). This was formulated as a quadratic programming problem and solved numerically.

**Figure 4.**
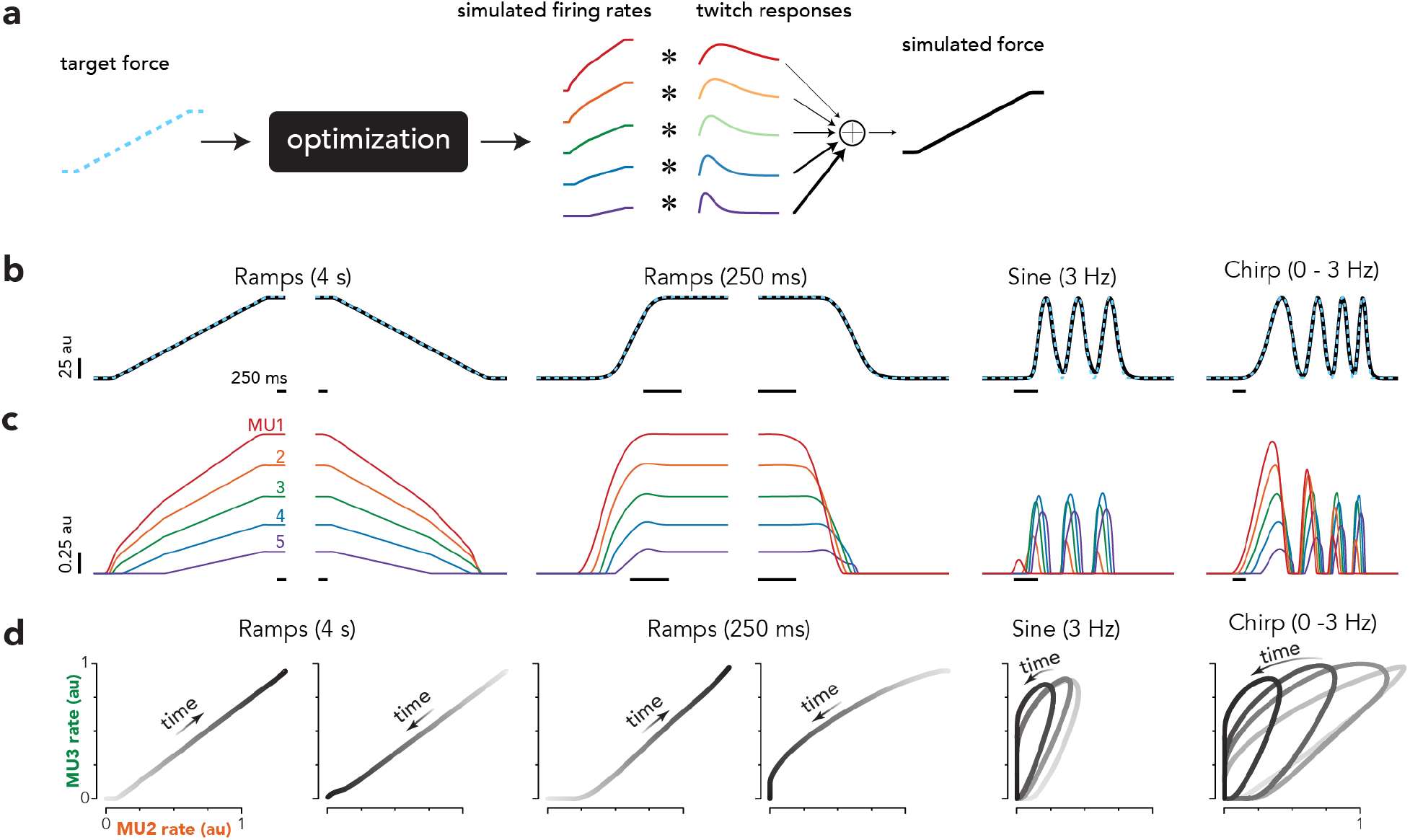
Optimal MU recruitment. (**a**) Isometric force production was modeled using an idealized MNP containing 5 MUs. MU twitch amplitude varied inversely with contraction time such that small MUs were also slow. The optimal set of MU firing rates for generating a target force profile was numerically derived as the solution that minimized the mean-squared error (including both mean error between the simulated MNP and target forces and variance of the simulated force). (**b**) Example target (*cyan*) and mean MNP (*black*) forces (for simplicity no variability is shown). (**c**) Optimal MU firing rates used to generate the MNP forces in ***b***. Each color corresponds to a different MU, numbered in ascending order by size (i.e., MU1 was the smallest and slowest). Optimization predicted size-based recruitment for steady force profiles (*first two columns*), but more flexible recruitment strategies for rapidly changing forces. (**d**) State space plot of MU3 (*green*) and MU2 (*orange*), for each condition shown in ***c***.

The model accurately generated force profiles (**Fig. 4b**) by leveraging a flexible interplay between MUs with different size and speed properties (**Fig. 4c**). Recruitment was purely size-based for slowly ramping forces. During faster ramps, deviations from purely size-based recruitment were modest and brief. If similar deviations were observed empirically, most would be indistinguishable from latency differences. However, recruitment became noticeably different during fast sinusoids. A similar effect was observed within chirps; recruitment changed as frequency increased. These solutions reflect the fact that, at low frequencies, using small (slow) MUs minimizes fluctuations^24,26^. At high-frequencies, it becomes optimal to rely more on large (fast) MUs to avoid excessive force during troughs. State-space plots confirm that, even under the assumption of complete flexibility, activity followed a similar manifold for all ramps (**Fig. 4d**). In contrast, flexibility revealed itself when comparing between the slow ramps and the high-frequency sinusoid, or when comparing within the chirp.

### Motor unit activity across force profiles

We recorded from the *deltoid* and *triceps* during the production of diverse force profiles (**Fig. 1d**) over 14 daily sessions (average of 347 trials per session and 29 per condition; range of 3-20 MUs per session, 134 total). MUs showed a range of recruitment thresholds and peak firing rates (**Fig. S2a**). There was no indication of general fatigue across a session (**Fig. S2b**). As above, analysis was restricted to simultaneously recorded neighboring MUs.

As predicted by the hypothesis of flexible control, MU recruitment differed between slowly changing forces and high-frequency sinusoids. **In Fig. 5c**, three *lateral triceps* MUs are active during the slow ramp (*left*). During the 3 Hz sinusoid, these MUs remain similarly (or slightly less) active, and are joined by a handful of MUs that are active primarily during the sinusoid (*right*). **Fig. S3** shows additional single-trial traces, and rasters for all trials / MUs from this session. Of 7 recorded MUs, three were strongly active during the 3 Hz sinusoid despite being inactive during the ramp.

**Figure 5.**
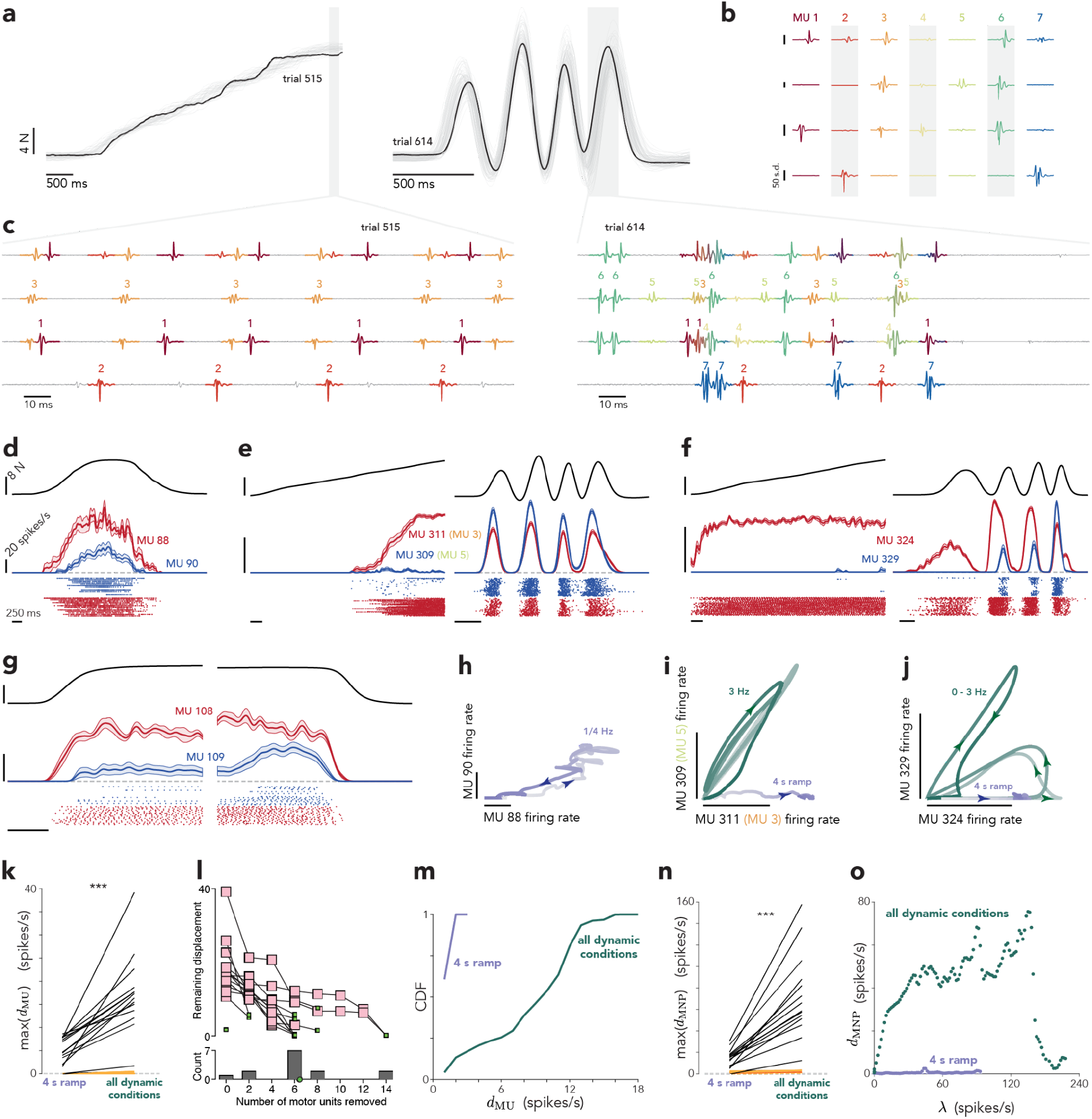
Dynamic experiments. **(a)** Forces for two conditions: four-second ramp 3 Hz sinusoid. Gray traces show all trials in this session. Black traces highlight trials for which EMG data are shown below. **(b)** MU spike templates across four (of eight) EMG channels recorded in the lateral triceps. Vertical scale: 50 standard deviations of the noise. Each template spans 10 ms. **(c)** EMG voltage traces for the two trials and time-periods highlighted in *a*. **(d,e,f)** Example MU responses. Each panel shows the trial-averaged force (*top*), mean firing rate with standard error (*middle*), and spike rasters (*bottom*). Vertical scales: 8N and 20 spikes/s. Horizontal scale: 250 ms. MU311 and MU309 correspond to MUs 3 and 5 in *c*. **(h,i,j)** State-space plots for the MUs shown in *d,e,f*. Scale bars: 20 spikes/s. (**k**) Maximum displacement for every session. Displacement was computed twice: within the 4-second ramp only, and again across all conditions. *Orange lines* plot displacement expected due to sampling error (ten resampled artificial populations per empirical dataset). Empirical slopes were significantly larger than expected given sampling error (two-sample t-test, p < 0.001, N=14 sessions). (**l**) Maximum displacement (across all conditions) after repeatedly removing the MU pair that caused the largest displacement. Each trace corresponds to one session, and ends when remaining displacement was no larger than the largest within the ten resampled populations (or when no pairs were left). Gray histogram plots the distribution of MUs removed (mean shown in *green*). One session (lone green square) terminated immediately because it contained only 3 MUs. (**m**) Cumulative density function for *d*_Mu_(*t*) for the slowly increasing ramp (*purple*) and all conditions (*green*) in one session. (**n**) As in *k* but for MNP dispersion. Slopes were significantly larger than expected given sampling error (two-sample t-test, p < 0.001, N=14 sessions). (**o**) *d*_MNP_(λ) as a function of the norm, λ, for the slowly increasing ramp (*purple*) and across all conditions (*green*) for one session.

Examples illustrate the range of effects. Activity tended to be consistent with rigid control during slowly changing forces (**Fig. 5d**) but could depart from rigid when comparing ramps with high-frequency sinusoids. In **Fig. 5e** MU309 becomes far more active during the fast sinusoid, even as MU311 becomes slightly less active (these MUs correspond to MU5 and MU3 from *panel c*). In **Fig. 5f**, MU329 is inactive during the slow ramp and early in the chirp. MU329’s rate then climbs for the higher-frequency chirp cycles, even as MU324’s rate declines. Consistent with both the optimal-recruitment model and prior experiments, fast ramps only rarely produced recruitment that was clearly incompatible with rigid control, despite the high rate-of-change (~64 N/s). Yet this did occasionally occur at force offset: MU109 tripled its activity in anticipation of a fast downward ramp, even as MU108 became slightly less active (**Fig. 5g**, *right*). A similar but smaller effect is exhibited by the model -- large/fast MUs become slightly more active, ‘taking over’ so force can terminate swiftly, a strategy that became more pronounced for faster offsets (not shown). A key aspect of this strategy is that it involves ‘looking ahead’ in time. Consistent with this interpretation, the increase in MU109’s rate reflected anticipation of force offset; no increase occurred when holding a static force over an extended time.

Direct inspection of individual-MU activity yields the opportunity to determine whether departures from rigid control might be indirectly due to other phenomena. Departures could not be explained by some MUs reflecting force while others reflected its derivative. MU309 (**Fig. 5e**) and MU329 (**Fig. 5f**) are more active during higher-frequency forces, but do not phase lead their neighboring MUs. The rate of MU109 (**Fig. 5g**) rises while force is constant. Rapidly increasing force ramps (e.g., **Fig. 5g**, *left*) did not cause rates to overshoot, which would occur if some MUs reflected the derivative. This hypothesis is further considered in model-based analyses below. Large departures from rigid control were also not explainable by some sort of rapid within-trial fatigue. There is no evidence of fatigue in **Fig. 5e**. In **Fig. 5f**, one might propose that MU324’s rate declines during the chirp due to fatigue, yet it held a consistently high rate during the much longer ramp. In **Fig. 5g**, the departure occurs because activity increases in anticipation of force offset. Still, some MUs did show a slight rate ‘sag’, especially during the long ramp (e.g., **Fig. 2e**, *left*). These long-timescale effects should presumably not be considered a meaningful departure from rigid control, and this is taken into account when interpreting quantitative analyses below. We occasionally observed fatigue-combatting rotations; an MU could become less active for a handful of trials. To reduce any condition-specific impact on trial-averaged rates, trials for all conditions were interleaved. Inspection of individual MU responses (e.g., **Fig S3**) also reveals that different recruitment between ramps and sinusoids cannot be attributed to rotations or other long-timescale fatigue-related effects.

In situations where activity was potentially consistent with rigid control (**Fig. 5d**), trial-averaged trajectories lay on a monotonic manifold (**Fig. 5h**) with only minor departures comparable to the SEM. When activity was incompatible with rigid control (**Fig. 5e,f**), trajectories diverged from a monotonic manifold (**Fig. 5i,j**). We quantified population-state trajectories via the displacement and dispersion metrics, concentrating on the prediction that departures from rigid control should be small during the slowly increasing ramp (where rigid and flexible control make the same predictions) but common when comparing across all conditions. The slowly increasing ramp is also a useful comparison because it evokes activity over four seconds, and thus provides an estimate of departures from idealized rigid control due to modest temporal instabilities. Indeed, during the ramp, both displacement (**Fig. 5k**) and dispersion (**Fig. 5n**) were small but higher than expected given sampling error (*orange lines*). (The latter was very low because displacements due sampling error were small and often correctable by the temporal shifts we allowed to eliminate latency-based effects). Inspection of individual cases revealed that the modest displacement during the slow ramp was primarily due to uninteresting phenomena such a slight sag in the rate of one MU over long timescales (as in **Fig. 2e**, *left*). We saw no compelling evidence for recruitment flexibility during slow ramps or other low frequency force profiles.

When considering all conditions, displacement (**Fig. 5k**) and dispersion (**Fig. 5n**) became much larger (p < 0.001, t-test comparing increase in slope for the data versus control for sampling error). All sessions had a maximum displacement outside the range expected due to sampling error. As above, we removed the MU pair with the largest oppositional change, and iterated until remaining displacement was within the range possible due to sampling error. On average, 6.4 (out of an average of 9.6) MUs had to be removed. Populations typically exhibited a range of large, medium and small displacements (**Fig. 5m**). The long-duration slow ramp provides an estimate of ‘trivial’ displacements due to temporal instabilities (*purple*). These were small. The distribution was shifted strongly to the right when considering all force profiles (*green*). For dispersion we assessed robustness by sweeping the norm, λ, used to compute dispersion (rather than taking the maximum). When considering all conditions (*green*), dispersion was high across a broad range of λ, indicating that population trajectories were far from hewing to a 1D monotonic manifold. In contrast, dispersion was always minimal when considering only the slow ramp (*purple*).

The contrast between the four-second ramp and sinusoidal forces also serves as a natural control for the concern that departures from rigid control might be due to occasional missed spikes. Judged subjectively, spike-sorting was excellent, but it remains likely that spikes were sometimes missed. For example, a small waveform might occasionally be missed if it overlaps with much larger waveforms when overall activity is high. Errors in spike sorting would, to a first approximation, be expected to impact all conditions similarly. Yet departures from rigid control were strongly present for some conditions but not others. One could argue that sorting might become particularly challenging during high-frequency sinusoids, leading to more missed spikes. However, the empirical violations of rigid control typically involved some MUs becoming more more active, not less active, during sinusoids. Direct inspection (**Fig. 5c**; **Fig. S3c**) of voltage traces also rules out this potential source of effects.

### Motor unit activity across muscle lengths

The solutions exhibited by the optimal-recruitment model suggest a broader phenomenon: if recruitment seeks optimality, it must depart from purely size-based whenever MUs show task-relevant diversity in domains other than size. To explore one potential domain, we employed a task with two postures. The *anterior deltoid* generated torque in the same axis (shoulder flexion) for both, but was at different lengths as it did so. We recorded MU population activity during six daily sessions (average of 495 trials per session; range of 4-9 MUs per session, 39 total). Postures could not be interleaved on individual trials, and were thus performed in blocks. The possibility of length-based flexibility has only rarely been considered^77^. A challenge is that slight changes in electrode location relative to muscle fibers can impact spike amplitude and complicate sorting^46,77^. Our recording and spike-sorting pipeline overcame this problem by relying on waveform shape across both time and many channels. The matched filter approach to spike-sorting^66^ was naturally insensitive to modest changes in amplitude, so long as the spatio-temporal profile was preserved. Example voltage traces are shown for two postures (**Fig. 6a,b**; only two channels shown for simplicity). Both MU3 and MU4 exhibit waveforms that shrink modestly from the first to second posture, yet could be accurately identified. MU5 was active only when the deltoid was shortened. More subtly, MU3 was slightly more active at the shorter length while MU4 was considerably less active. This can be seen by counting spikes in the voltage traces, but is more readily observed in **Fig. 6e** (*top-right subplot;* session-specific identifiers MU3 and MU4 correspond to global identifiers MU157 and MU158).

**Figure 6.**
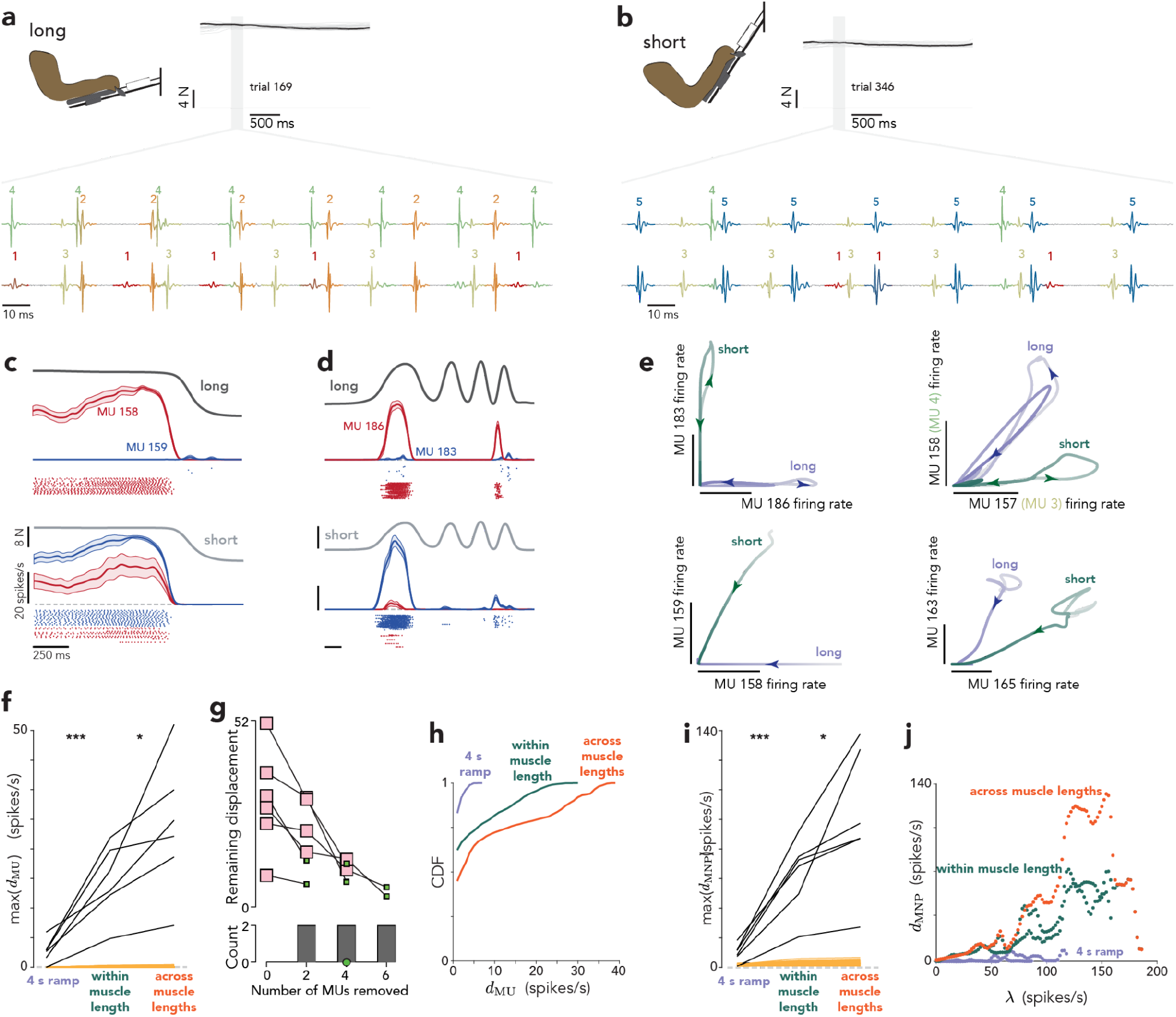
Muscle-length experiments. The monkey generated a subset of the forces in *Fig. 1d* with a shoulder-flexion angle of 15° (the deltoid was long) or 50° (the deltoid was short). **(a)** Force as a function of time (*gray traces*) for all trials for one condition (static force, long posture) in one session. EMG voltages traces are shown for the highlighted portion (*gray window*) of an example trial (*black*). **(b)** Same as in *a* but for the short posture. **(c, d)** Example MU responses. Each panel shows the trial-averaged force (*top*), mean firing rate with standard error (*middle*), and spike rasters (*bottom*). Vertical scales: 8N and 20 spikes/s. Horizontal scale: 250 ms. (**e**) State space plots for the MUs shown in *c,d* and two additional example pairs. MUs 157 and 158 correspond to MUs 3 and 4 in *a,b*. **(f)** Maximum displacement for every session, computed within the 4-second ramp, across all conditions within a posture, and across all conditions for both postures. *Orange lines* plot displacement expected due to sampling error (ten resampled artificial populations per empirical dataset). *** and * symbols indicate p<0.001 and p<0.05 respectively (two-sample t-test, N=6 sessions). (**g**) Maximum displacement (across all conditions) after repeatedly removing the MU pair that caused the largest displacement. Each trace corresponds to one session, and ends when remaining displacement was no larger than the largest within the ten resampled populations. Gray histogram plots the distribution of MUs removed (mean shown in *green*). (**h**) Cumulative density function for *d*_MU_(*t*) for the slowly increasing ramp (*purple*), across conditions within a posture and then pooled across postures (*green*), and across all conditions for both postures (*orange*), for one session. (**i**) As in *f* but for MNP dispersion. (**k**) *d*_MNP_(λ) as a function of the norm, λ, for one session.

Muscle-length-driven changes in recruitment (**Fig. 6c,d**), yielded state-space trajectories incompatible with a single monotonic manifold (**Fig. 6e**). This produced high values of displacement (**Fig. 6f**) and dispersion (**Fig. 6i**). Because recruitment differed across force profiles, displacement and dispersion were high even when computed within a muscle length (*green*). They became higher still when computed across muscle lengths (*orange*) due to length-specific recruitment. All sessions had maximum displacement greater than expected due to sampling error. On average, more than half the MUs (4 out of an average of 6.5 recorded MUs) had to be removed from the population to eliminate significant displacement. The overall distribution of displacements (**Fig. 6h**, *orange*) was shifted strongly rightwards relative to that observed during the slow ramp at one muscle length (*purple*). Dispersion was high across a broad range of values of the norm (λ) at which it was evaluated (**Fig. 6j**), due to the very different trajectories for different muscle lengths as illustrated in panel *e*.

Effects were unrelated to fatigue over the course of the daily session. Recruitment changed suddenly at block boundaries, not slowly over the course of the session (e.g., *blue rasters* in **Fig. 6d** display a sudden increase between *top* and *bottom* subpanels, not a gradual change spanning both). A related possibility is that the near-complete inactivity of a previously active MU might be due to recording instabilities. This is unlikely; other MUs recorded on the same channel remained visible, indicating there was minimal electrode movement. Furthermore, often an MU became much less active but still spiked, ruling out this concern (**Fig. S4**). Additionally, in some sessions we re-examined the first posture, and confirmed that an MU that fell inactive during the second posture became active upon returning to the first posture.

### Latent factor model

The most straightforward test of a hypothesis is to formalize it as a model and ask whether it can fit the data. With the hypothesis of rigid control, this has historically been challenging because of the large space of free parameters. Existing models of rigid control use constrained link functions (rectified linear^73^ or sigmoidal^74^). These are reasonable but limit expressivity. The ideal model should have only those constraints inherent to rigid control – only then can it be rejected if it fails to fit the data. Fortunately, machine-learning methods now allow fitting models with unconstrained link functions and unknown common drive. By fitting many simultaneously recorded MUs across diverse conditions, one can obtain a stringent test of rigid control despite the large free-parameter space.

We employed a probabilistic latent factor model (**Fig. 7a**) where each MU’s rate is a function of common drive: *r_i_*(*t*) ~ *f_i_*(*x*(*t* + *τ_i_*)) Fitting used black box variational inference^78^ to infer *x*(*t*) and learn the MU-specific *f_i_* and time-lag, *τ_i_. f_i_* was unconstrained other than being monotonically increasing. The resulting model can assume essentially any common drive and set of link functions, and MUs can have different latencies.

**Figure 7.**
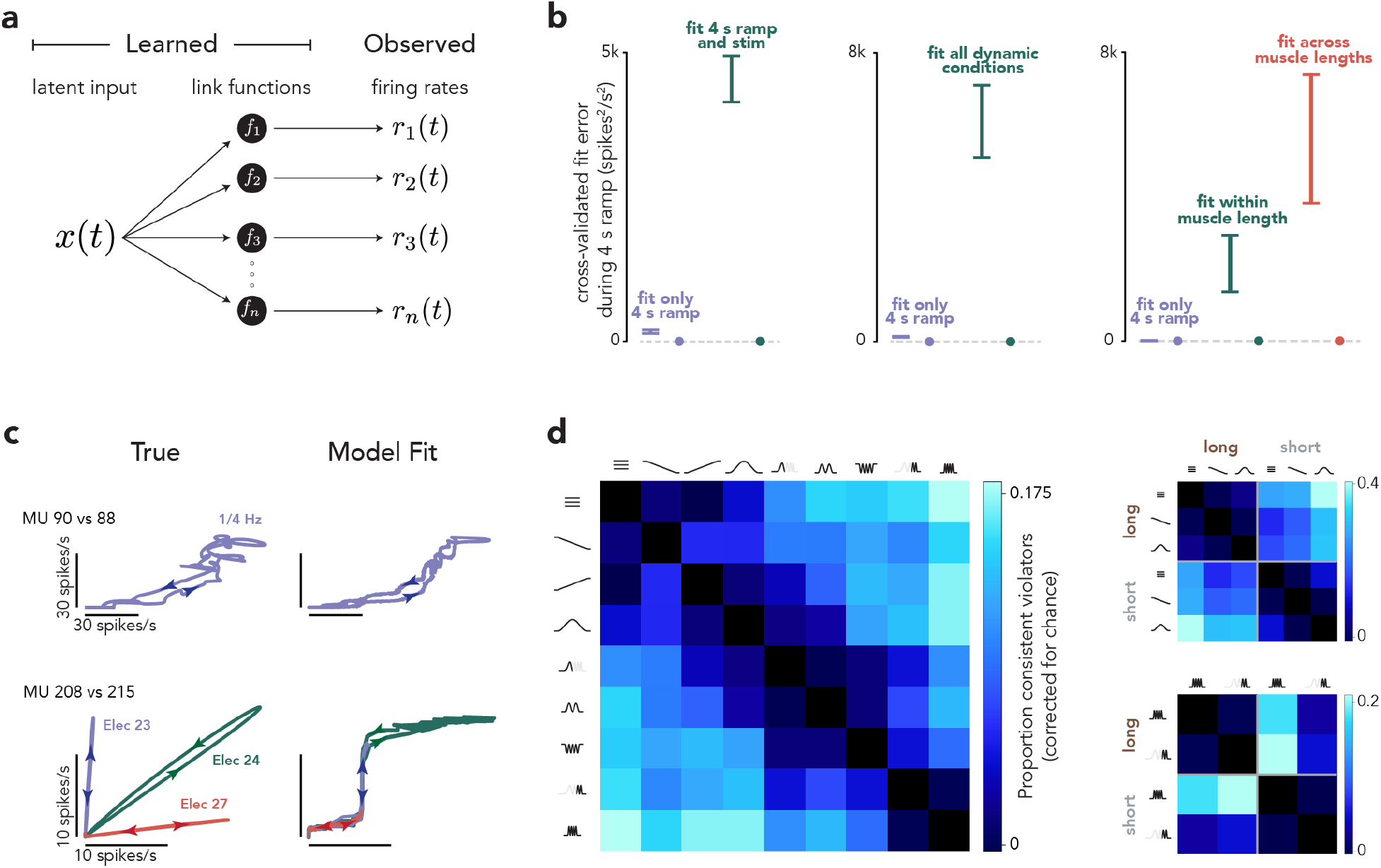
Latent factor model. (**a**) The model embodies the premise of rigid control: MU firing rates are fixed ‘link functions’ of a shared, 1-D latent input. (**b**) Model performance for stimulation (*left*), dynamic (*middle*) and muscle-length (*right*) experiments. The model was fit separately for each session. Regardless of how many conditions were fit, cross-validated error was computed only during the four-second ramp. Cross-validated error involved fitting separately to two data partitions. Partitioning was repeated 20 times. Each time we randomly partitioned the trials for each session, fit the model, then computed the overall fit error as the median across all MUs in all sessions. Vertical bars indicate the mean and standard deviation across these 20 partitionings, and indicate reliability with respect to sampling error. *Purple bars* plot error when fitting only the four-second ramp. For stimulation experiments (*left*), *green bars* plot error when also fitting conditions where stimulation was delivered. For dynamic experiments (*middle), green bars* plot error when fitting all force profiles. For muscle-length experiments (*right*), *green bars* plot error when fitting all force profiles for the same muscle length (then averaging across muscle lengths). *Orange bars* plot error when fitting all conditions. Circles show cross-validated fit error when for an artificial population response that obeyed rigid control, but otherwise closely resembled the empirical population response. (**c**) Illustration of model fits for simplified situations: the activity of two MUs during a slow ramp (*top*) or following cortical stimulation on three different electrodes (*bottom*). For illustration, the model was fit only to the data shown. (**d**) Proportion of total MUs that consistently violated the single-latent model when fit to pairs of conditions. Each entry is the difference between the proportion of consistent violators obtained from the data and the proportion expected by chance. *Left*: dynamic experiments. *Right-top*: muscle-length experiments. For muscle length experiments, we grouped conditions (by frequency) to focus on the impact of muscle length. *Top*, lower-frequency forces. *Bottom*, higher-frequency forces.

To be conservative, we ask whether the model can fit the data when given every opportunity, rather than testing generalization to held-out data. Some fit error is inevitable due to sampling error from spiking variability and other forms of trial-to-trial variability^79^. We thus evaluated whether the model made consistent, systematic errors. We fit the model to two random data partitions and computed cross-validated residual error based on the dot product of the two residuals^80^ (residuals will be uncorrelated if due to sampling error). Cross-validated error can be positive or negative, and should be zero on average for an accurate model and greater than zero for an inaccurate model. To confirm that optimization was successful we used two different methods to construct artificial datasets that obeyed rigid control but were otherwise realistic and contained sampling error (see Methods and **Fig. S5**). The latent model always fit artificial datasets successfully, with cross-validated error near zero (e.g., **Fig. 7b**, *filled circles*).

As before, we focused on a central prediction of optimal recruitment: rigid control should account for recruitment during slowly changing forces (where flexible and rigid control make the same predictions) but not across all conditions. For each session, we fit MU population activity during the four-second increasing ramp alone, and again during all conditions. Error was always computed only for the fit within the four-second ramp. A model embodying rigid control should fit this situation very well, unless it has to also fit other situations (which may require compromises because the same link functions may not work). The strategy of comparing error for the same condition is conservative, and avoids the concern that different conditions might be intrinsically easier or harder to fit.

The model performed well when fitting only the four-second increasing ramp. Cross-validated error was consistently above zero, but only very slightly (**Fig. 7b**, *purple*). This confirms that rigid control is an excellent model of MU recruitment during slowly changing forces, in agreement with the vast literature on this subject. Model failures due to trivial phenomena – e.g., short-timescale fatigue – are small.

Error rose dramatically when the model had to also account for other conditions (**Fig. 7b**, *green and orange*, results compiled across all sessions). The range of errors when fitting all conditions was much higher than the range when fitting only the ramp. On individual sessions, error ranges were non-overlapping for 13 of 17 stimulation experiments, 12 of 14 dynamic experiments, and 5 of 6 muscle-length experiments, (the analysis could not be performed for one stimulation experiment because there was little activity during the slow ramp). Failures were readily understood: the model could fit activity that followed a monotonic manifold during slowly changing forces (**Fig. 7c**, *top*), but not activity that lay on different manifolds across conditions (*bottom*).

Comparing only slowly and rapidly increasing ramps did not yield compelling evidence for flexibility. When asking the model to fit the rapidly increasing ramp in addition to the slowly increasing ramp, the error increase was modest: only 9% as much as when asking the model to fit all conditions. In contrast, the error increase was large (65% as much) when asking the model to fit the 3 Hz sinusoid in addition to the slowly increasing ramp.

This finding agrees with both the optimal recruitment model (**Fig. 4**) and what can be seen in examples: departures from rigid control are most prevalent when comparing forces that evolve continuously but have different frequency content. To explore further, we considered conditions that spanned a range of frequencies, from static to 3 Hz sinusoid (dividing the chirp into two sub-conditions). We fit the model of rigid control to single-trial responses for just two conditions at a time. We defined an MU as a ‘consistent violator’ if its overall activity was overestimated for trials from one condition and underestimated for trials from the other, at a rate much higher than expected by chance (computed using binomial statistics). Of course, even when a population response is overall inconsistent with rigid control, only some individual MUs will show effects large enough to be defined as consistent violators (especially when comparing only two conditions and ignoring within-condition violations). This is acceptable because we wish not to interpret absolute scale, but to assess which conditions tend to evoke different recruitment. Consistent violators were rare when two conditions had similar frequency content (**Fig. 7d**, *left, dark entries near diagonal*), and prevalent when conditions had dissimilar frequency content. For example, the hypothesis of rigid control failed most often when comparing between fast sinusoidal forces (the 3-Hz sinusoid and the end of the chirp) and slowly changing forces (the collection of three static forces, slowly increasing and decreasing ramps, and 0.25 Hz sinusoid).

Thus, MU responses don’t simply randomly depart from rigid control, but do so in ways expected under the hypothesis of flexible control. Violations of rigid control during muscle-length experiments were also systematic. We divided conditions into slowly changing forces (**Fig. 7d**, *top-right*) and quickly changing forces (*bottom-right*). Within each category, consistent violators were uncommon when comparing conditions with the same muscle length (dark block-diagonal entries), and more common when comparing across muscle lengths.

The model of rigid control we applied is unusually expressive. Yet might it be possible to increase expressivity, and perhaps achieve better fits, while still maintaining the core assumption of common drive? Further increasing the expressivity of the link functions with more free parameters had almost no effect. We also employed a model that allowed history dependence: there was a single common drive, but each MU could uniquely reflect the current value of that drive and/or integrate past values. This allowed the model to display features not normally considered part of rigid control. For example, the inferred common drive could reflect force, and some MUs could be directly sensitive to that force and others to the history of force. Alternatively, the inferred common drive could reflect a high-passed version of force, allowing some MUs to reflect the rate-of-change of force while others (via low-pass filtering) reflect force. Fits were improved for some small specific features likely caused by persistent inward currents. Yet there was still a dramatic increase in error when the model had to account for responses during all conditions.

In contrast, excellent fits could be obtained by abandoning the core assumption of common drive and allowing multiple latent factors (multiple ‘drives’), as proposed by flexible control. To ask how many latent factors might be needed (**Fig. 8a**), we challenged the model using the two dynamic sessions with the most simultaneously recorded MUs (16 and 18 recorded from the *triceps*, selecting for sessions where all MUs were reasonably active at some point). Expanding the number of drives produced an immediate decrease in cross-validated error, which reached zero around 4-6 factors (**Fig. 8b**). Thus, the model can fit well, it simply needs more drives than allowed under rigid control.

**Figure 8.**
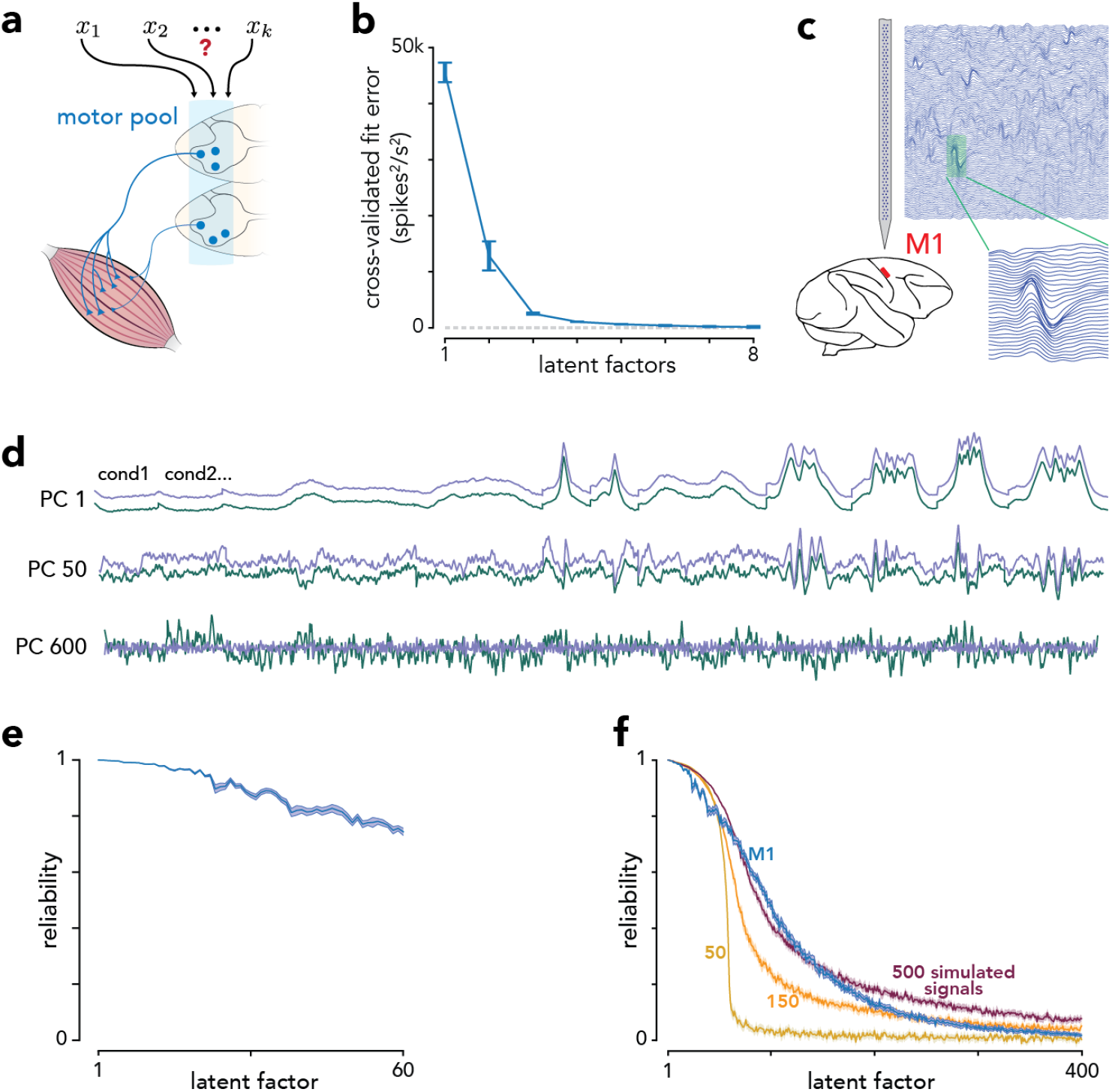
Quantifying neural degrees of freedom. (**a**) Schematic illustrating the central question: how many degrees of freedom might influence the MNP? (**b**) Cross-validated fit error for all conditions when fitting two dynamic-experiment sessions using models with 1-8 latent factors. Error bars indicate the mean +/- standard deviation of that median error across 10 model fits, each using a different random partition to compute cross-validated error. (**c**) We recorded neural activity in M1 using 128-channel Neuropixels probes. (**d**) Two sets of trial-averaged rates were created from a random partitioning of trials. Traces show the projection of the first (*green*) and second (*purple*) onto PCs found using only the first. Traces for PCs 1 and 50 were manually offset to aid visual comparison. (**e**) Reliability of neural latent factors for the first 60 factors. Traces plot the mean and central 95% (*shading*) of the distribution across 25 random partitionings. (**e**) Same but latent factors 1-400. The empirical reliability (*blue*) is compared to that for simulated data with 50 (*yellow*), 150 (*orange*), and 500 (*red*) latent signals.

### Neural degrees of freedom

We recorded a minority of MUs (the *triceps* alone contain a few hundred) during a subset of behaviors and found that describing activity required 4-6 drives. Neural control of the arm may thus be quite high-dimensional, with dozens or even hundreds of degrees of freedom across all muscles. It is unclear what proportion of these are influenced by descending control, but it could be many. The corticospinal tract alone contains approximately one million axons in human^81^, and our stimulation experiments demonstrate remarkably fine-grained control.

For flexible descending control to be plausible, motor cortex activity would need to display a great many degrees of freedom. However, motor cortex activity is often described as being constrained to relatively few degrees of freedom^82–84^, with 10-20 latent factors accounting for most response variance^83–87^. Studies have stressed the presence of more degrees of freedom than predicted by simple models^88,89^, but even the largest present estimate (~30^88^) seems insufficient to support extensive flexible control over many muscles. However, the meaning of these numbers is unclear and they depend on methodology. For example, in^88^, a variance-explained threshold yielded an estimated dimensionality of 9, while assessing signal reliability yielded an estimate of 29. Both are likely underestimates given analyzed populations on the order of 100 neurons. The present task yields the opportunity to revisit this issue using a task relevant to flexible control. We considered an unusually large population (881 sulcal neurons) recorded over 13 sessions (22 - 128 neurons per session) using the 128-channel version of primate Neuropixels probes.

Capturing 85% of the variance required 33 dimensions. This is high relative to prior estimates that used a variance cutoff, possibly due to condition variety in the Pac-Man task. Yet we stress that variance explained is not a particularly meaningful measure given the scientific questions at hand. Small-variance signals may be very relevant to descending control^90^. Thus, when asking whether a latent factor is meaningful, we focused not on size but on whether the factor was reliable across trials (**Fig. 8d**) using a method similar to that of Stringer and colleagues^91^. We defined reliability, for the projection onto a given principal component, as the correlation between held-out data and the data used to identify the principal component. Even though each PC past the first 33 captured a small signal, those signals were reliable (**Fig. 8e**); the first 60 all had reliability >0.7.

Reliability remained well above zero for approximately 200 PCs (**Fig. 8f**). To provide context, we analyzed artificial datasets that closely matched the real data but had known dimensionality. Even when endowed with 150 latent factors, artificial populations displayed reliability that fell faster than for the data. This is consistent with the empirical population having more than 150 degrees of freedom, an order of magnitude greater than previously considered^83–87^. For comparison, if M1 simply encoded a force vector, there would be only one degree of freedom; all forces in our experiment were in one direction. Encoding of the force derivative would add only one further degree of freedom. Thus, the M1 population response has enough complexity that it could, in principle, encode a great many outgoing commands beyond force *per se*. Direct inspection of individual-neuron responses supports this view; neurons displayed a great variety of response patterns (**Fig. S6**).

## Discussion

The hypothesis of rigid control has remained dominant^13,14^ for three reasons: it describes activity during steady force production^16,72,75,92–98^, would be optimal in that situation^24^, and could be implemented via simple mechanisms^19,21^. It has been argued that truly flexible control would be difficult to implement and that “it is not obvious… that a more flexible, selective system would offer any advantages.^21^” Our findings argue that flexible MU control is a normal aspect of skilled performance in the primate. Recruitment differed anytime two movements involved different force frequencies or muscle lengths.

Frequency selective recruitment supports the proposal that it could be beneficial to use different recruitment strategies when modulating force slowly versus quickly^51^. Skeletal muscle fibers are functionally and structurally heterogeneous, yielding diversity in activation and relaxation times across MUs^3,8^. Activation-deactivation dynamics impacts the mechanical work produced by a muscle during rapid cyclic contractions, which has been suggested to warrant speed-based neural control mechanisms^99^. It is known that different muscles, with different compositions of slow- and fast-twitch fibers, become coordinated differently during rapid paw shakes^100^, bust-glide swimming^101^, and cycling^102,103^. Yet it has remained controversial whether similar differences might occur at the level of individual MUs within a muscle^44,56^. Most studies of single MUs (e.g., ^36,52,53,57^) report no speed-based change in recruitment beyond trivial effects of latency. Thus, the prevailing view is that speed-based recruitment either doesn’t occur^13,44,55^, or occurs only in a subset of lengthening contractions^56^.

A factor underlying this controversy may be the different force profiles used in different studies. The failure to find speed-based recruitment during step-like force profiles is consistent with our simulations. Even with complete flexibility, the optimal solution during a sudden force increase is to largely maintain the pattern of recruitment used for slowly increasing forces. In contrast, the optimal-recruitment model predicts that recruitment should become quite different during high-frequency sinusoids and chirps. Consistent with this, the few studies that have argued for speed-based recruitment flexibility involved sinusoidal forces^10,48–50^. Prior evidence was indirect (based on the EMG spectrum, where interpretation can be ambiguous^63^) or limited (e.g., a single example where two MUs were recorded simultaneously.) Yet our data argue that the conclusion of these studies was correct. We found it was common for some MUs to be relatively inactive during slowly changing forces but to become strongly active during fast sinusoids or at the end of chirps.

A simple way of achieving very limited frequency-based flexibility would be to allow some MUs to reflect force and others to reflect rate-of-change of force. This was not the strategy used by the optimal-recruitment model, but might it explain the empirical data? This is unlikely. First, greater responsivity during high frequencies did not imply phase shifted responses (e.g., earlier peaks) as would be predicted by sensitivity to rate-of-change. Second, rapidly increasing force ramps rarely led to compelling departures from rigid control. Finally, when fitting the data with multiple latent factors, it was not the case that one drive was the derivative of the other.

We observed MUs that became less active at the highest frequencies (e.g., **Fig. 5f**) but never observed an MU that became inactive at high frequencies, even though the optimal-recruitment model predicted this should occasionally happen. This discrepancy may reflect the model’s simplicity: it doesn’t take into account that slow fibers can speed up under negative load^104^, or may need to be shortened even when not contributing force. Yet despite being overly simple, the optimal-recruitment model is sufficient to make the key high-level point: accurately generating force profiles while minimizing other costs will typically require respecting diversity not only of size, but in any domain relevant to the range of behaviors. Consistent with this, muscle length also had a large impact on MU recruitment. Improved recording technologies, such as high-density multi-channel EMG electrode arrays^105,106^, are poised to allow future exploration of other domains^107^.

Recruitment flexibility likely reflects both spinal and supraspinal mechanisms. In the present case, muscle-length-driven flexibility presumably depends upon spinally available proprioceptive feedback^108^. During dynamic movements, some aspects of flexibility reflect future changes in force, knowledge of which presumably requires descending signals. The nature of the interplay between spinal and descending contributions remains unclear, as is the best way to model flexibility. Flexibility could reflect multiple additive drives to the MU pool and/or modulatory inputs that alter input-output relationships^71,109^ (i.e., flexible link functions). Both mechanisms could have spinal sources (changes in joint angle alters the intrinsic properties of motoneurons^110^) and/or supraspinal sources (different MU types receive dissimilar patterns of synaptic input via the pyramidal tract^68^).

The hypothesis that descending signals can influence MU recruitment has historically been considered implausible, as control might be unmanageably complex unless degrees of freedom are limited^21,28^. Indeed, descending control has typically been considered to involve muscle synergies^111^, above rather than below the level of individual muscles. This view accorded with the finding that stimulation of various brain regions evoked, with rare exceptions^69^, responses consistent with size-based recruitment ^12,35^. Yet we found that highly selective recruitment was readily observed from cortex. Even before insertion of EMG electrodes, we sometimes observed that a given cortical site activated only a subset of MUs within a muscle, forming a discrete spatial patch of visible contraction. It remains unclear whether this selective activation occurs at the level of small or large groups of MUs, but it is clearly well below the level of the muscles. Cortical control could potentially be more fine-grained in primates than in cats^81^, where most prior experiments were performed. If so, the evidence for selective recruitment in older studies may have been weaker and easy to dismiss as resulting from a damaged preparation^35^.

Our findings are potentially consistent with the observation that recruitment can be altered using biofeedback training in humans^47,58–61,112^. An open question has been whether subjects learn to exploit the fact that rigid control embodies (fixed) exceptions to pure size-based recruitment^46,62^, or whether subjects gain direct voluntary control over previously unknown degrees of freedom. Our results indicate the latter is possible but don’t directly address this question; the prevalence of flexibility during voluntary movement doesn’t necessarily imply it can be controlled under biofeedback.

A substantial literature argues that neural activity in most brain regions occupies far fewer dimensions than the number of recorded neurons^113^. This is true in motor cortex in an operational sense: a handful of high-variance signals^83–87^ are often informative when testing or developing hypotheses^114^. Yet there also exist small-variance signals that are reliable^88,90^ and thus could contribute to outgoing commands. The present results reveal that small-but-reliable signals are numerous. Although it is typically proposed that descending control of voluntary movement is and should be simple^13,115^, in our view there is little reason to doubt either the utility or capacity for controlling many degrees of freedom. The corticospinal tract alone contains on the order of a million axons^81^, including direct connections onto α-motoneurons^81,116^. Future experiments will need to further explore the level of granularity of descending commands, and how those commands interact with spinal computations.

## Acknowledgements

We thank Y. Pavlova for fantastic animal care. This work was supported by the Grossman Charitable Trust, Simons Foundation (M.M.C., J.P.C., and L.F.A.), McKnight Foundation (M.M.C. and J.P.C.), NIH Director’s DP2 NS083037 (M.M.C.), NIH CRCNS R01NS100066 (M.M.C. and J.P.C.), NIH 1U19NS104649 (M.M.C., L.F.A., and J.P.C.), NIH F31 NS110201 (N.J.M.), NIH K99 NS119787 (J.I.G.), NSF GRFP (N.J.M.), NSF NeuroNex (J.I.G. and L.F.A.), Kavli Foundation (M.M.C. and L.F.A.), Howard Hughes Medical Institute (M.N.S.), and Gatsby Charitable Foundation (J.I.G., L.F.A., and J.P.C.).

## Author Contributions

M.M.C. conceived the study; N.J.M., M.M.C., and S.M.P. designed experiments; N.J.M. collected and analyzed EMG data sets; N.J.M. and E.M.T. collected neural data sets, supervised by M.M.C. and M.N.S.; J.P.C and J.I.G. developed and trained latent factor models with input from M.M.C and N.J.M; E.A.A. and N.J.M. analyzed neural data sets; N.J.M. developed the optimal recruitment model, supervised by L.F.A. and M.M.C.; N.J.M. and M.M.C. wrote the paper. All authors contributed to editing.

## Code and Data Availability

The code and data that support the findings of this study are available from the corresponding author upon reasonable request.

---

**Figure S1.**
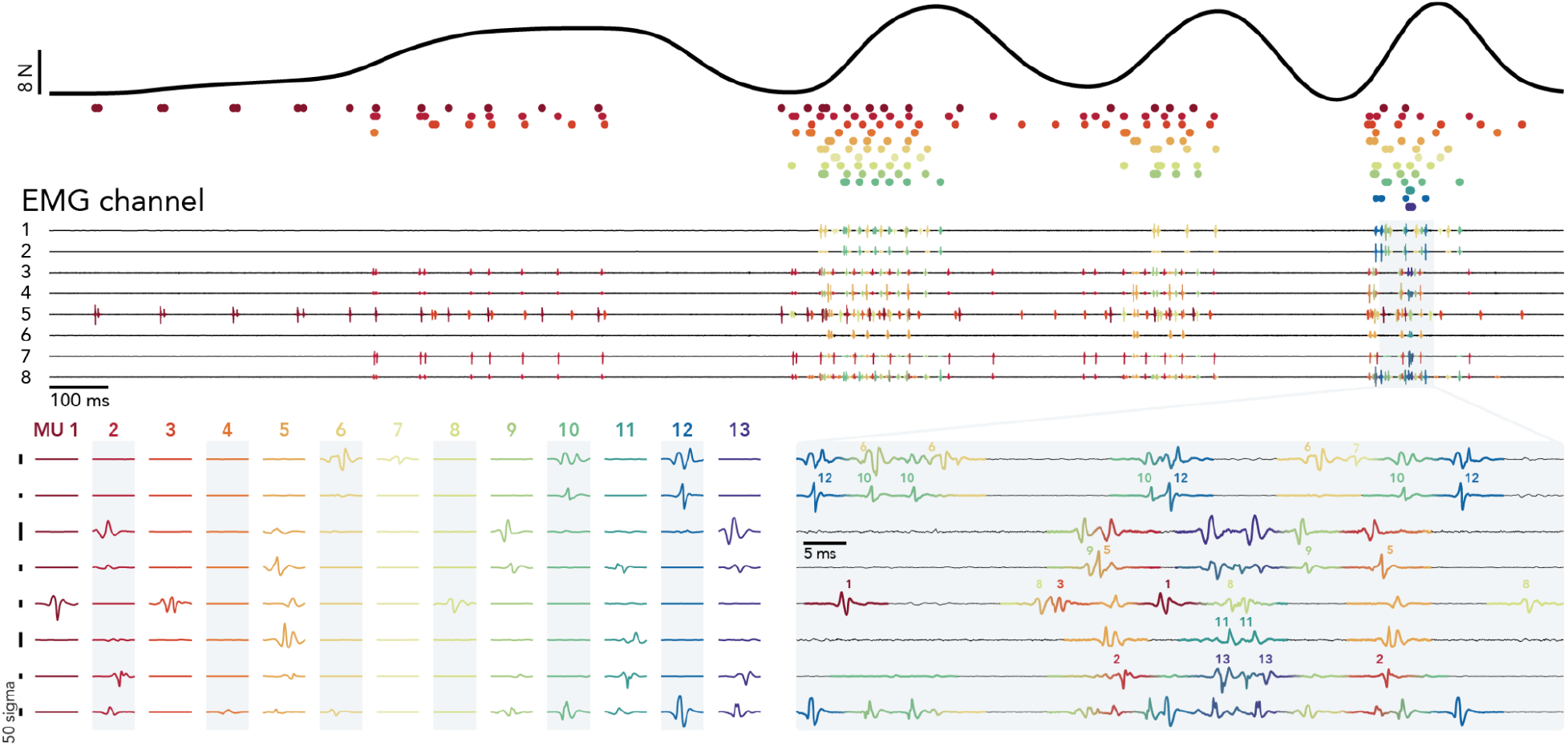
Example MU spikes. Behavior and MU responses during one dynamic-experiment trial. The target force profile was a chirp. *Top*: generated force. *Middle*: eight-channel EMG signals recorded from the lateral triceps. 20 MUs were isolated across the full session; 13 MUs were active during the displayed trial. MU spike times are plotted as *circles* (one row and color per MU) below the force trace. EMG traces are colored by the inferred contribution from each MU (since spikes could overlap, more than one MU could contribute at a time). *Bottom left*: waveform template for each MU (*columns*) and channel (*rows*). Templates are 5 ms long. As shown on an expanded scale (*bottom right*), EMG signals were decomposed into superpositions of individual-MU waveform templates. The use of multiple channels was critical to sorting during challenging moments such as the one illustrated in the expanded scale. For example, MU2, MU5, and MU10 had very different across-channel profiles. This allowed them to be identified when, near the end of the record, their spikes coincided just before the final spike of MU12. The ability to decompose voltages into a sum of waveforms also allowed sorting of two spikes that overlapped on the same channel (e.g., when the first spike of MU6 overlaps with that of MU10, or when the first spike of MU9 overlaps with that of MU5). The fact that sorting focused on waveform shape across time and channels (rather than primarily on amplitude) guarded against mistakenly sorting one unit as two if the waveform scaled modestly across repeated spikes (as occurred for a modest subset of MUs).

**Figure S2.**
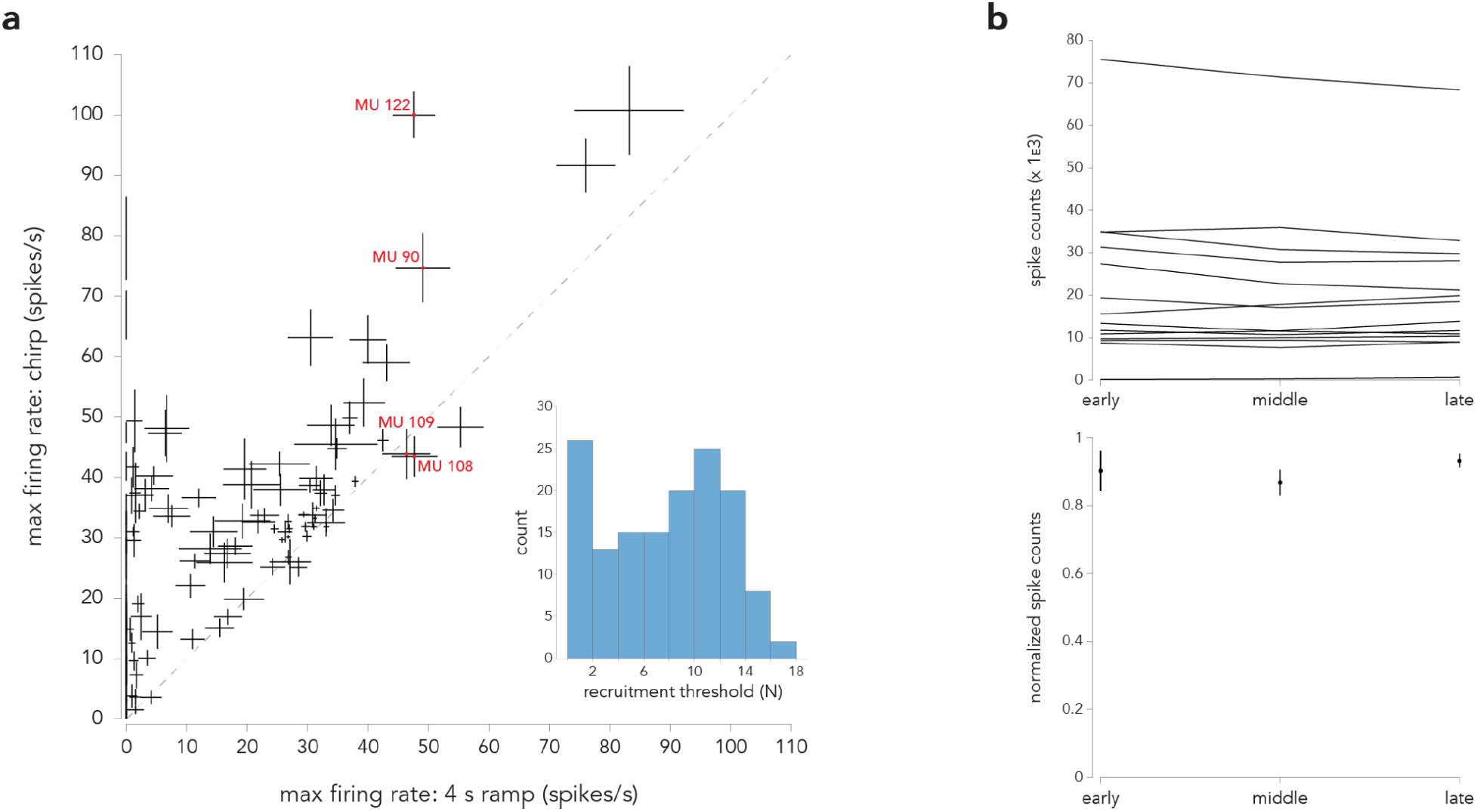
Basic properties of MU responses. (**a**) Comparison, for all MUs recorded during the dynamic experiments, of maximum rates during the four-second increasing ramp and during the chirp. Each point plots the maximum trial-averaged firing rate and its standard error for one MU. Labels highlight four MUs that have similar maximum firing rates during the ramp but different maximum firing rates during the chirp. There were also many MUs that were nearly silent during the ramp but achieved high rates during the chirp (points clustered near the vertical axis). *Inset*: distribution of recruitment thresholds, estimated as the force at which the MU’s firing rate, during the four-second ramp condition, exceeded 10% of its maximum rate during that condition. (**b**) Analysis of the possible impact of fatigue across the course of each session. *Top*: total MU spike counts, over the full recorded population, after dividing the session into thirds. Each line corresponds to one session. *Bottom*: Mean (across sessions) and standard error of the normalized total MU spike counts (counts for each session were normalized by maximum across trial epochs). There is little overall change in MU activity over the course of a session.

**Figure S3.**
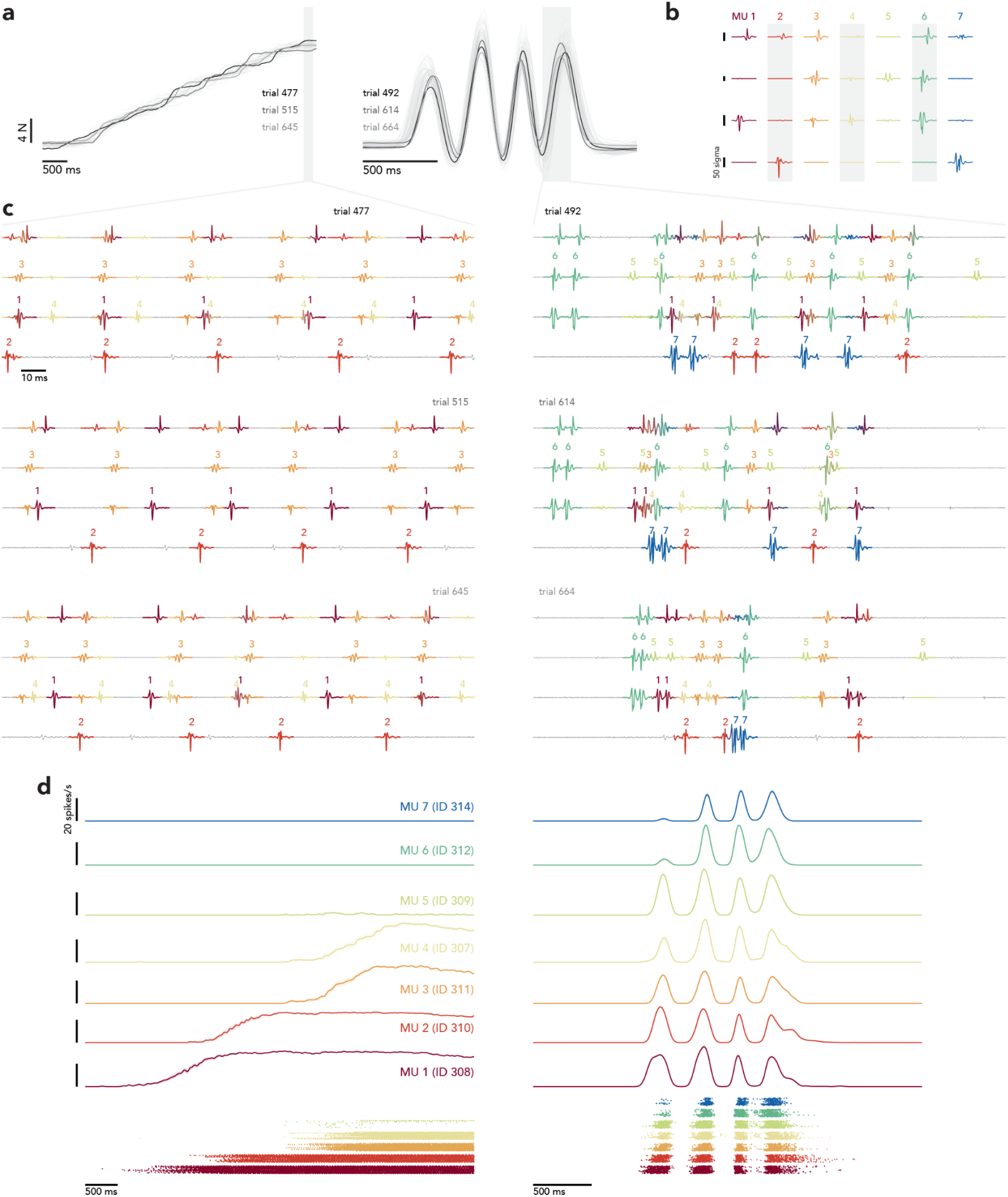
Additional documentation of single-trial responses during a dynamic experiment session. Presentation is similar to **Fig. 5a-c** but voltage traces are shown for six total trials, and spike rasters are shown for all MUs and all trials for two conditions (the four-second ramp and the 3 Hz sinusoid). (**a**) Same as **Fig. 5a** but traces are highlighted for six trials (three trials per condition) each of which is shown on an expanded scale below. (**b**) MU spike templates, repeated from **Fig. 5b**. (**c**) EMG signals for the highlighted trials and times. Six trials are shown in three rows (four channels are shown per trial). *Left column*: three trials for the ramp, the first of which repeats the trial shown in **Fig. 5c**. *Right column*: three trials for the sinusoid, the first of which repeats the trial shown in **Fig. 5c**. Vertical and horizontal scales as in ***b***. (**d**) Trial-averaged firing rates for each MU and condition (*top*) and spike rasters across trials (*bottom*). Vertical scale: 20 spikes/s. Horizontal scale: 500 ms. Spike rasters contain one row per trial, ordered from the first trial for that condition at the bottom to the last at top. Trials for all conditions were interleaved during the experiment.

**Figure S4.**
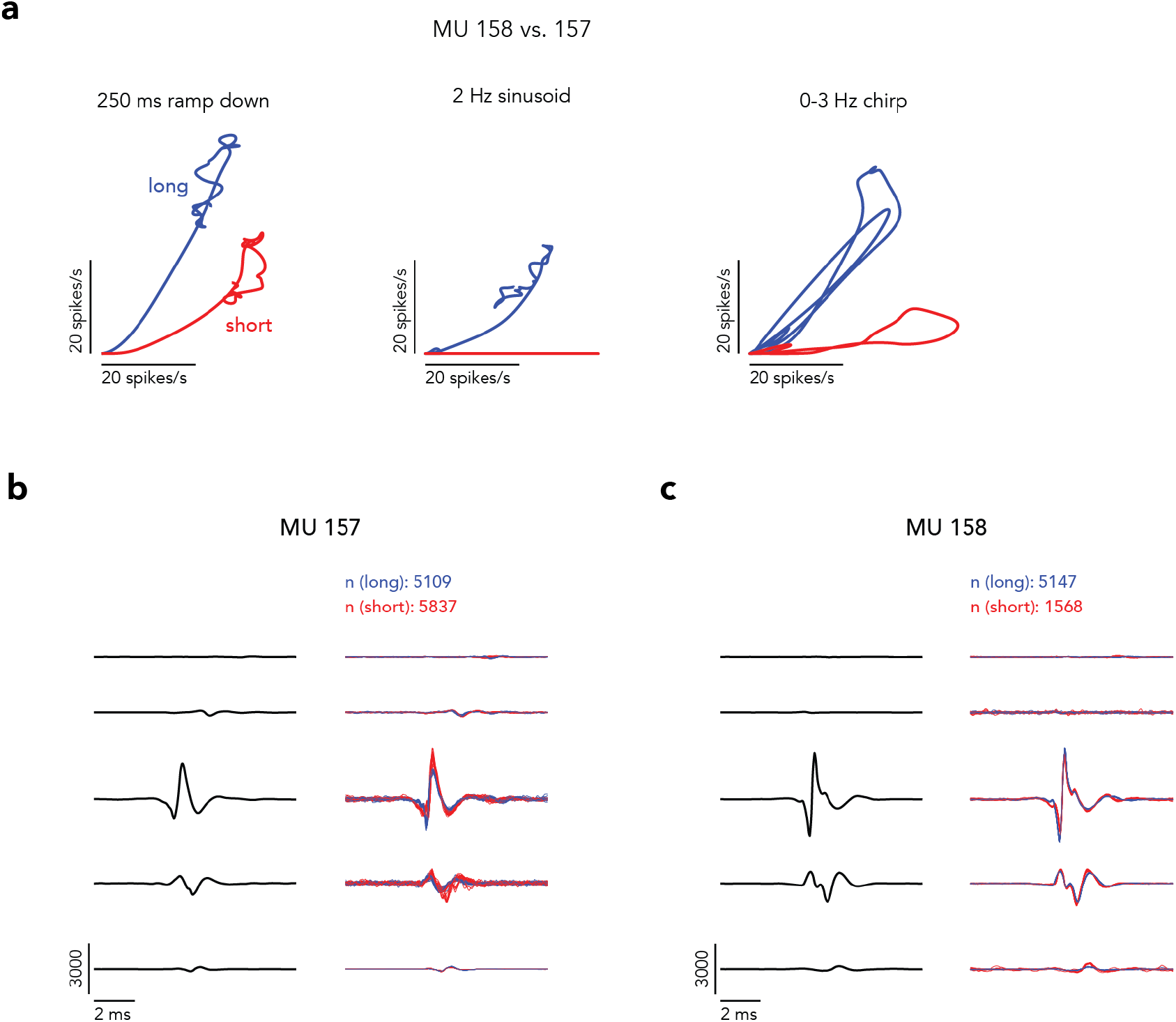
Example MU responses and waveforms across muscle lengths. (**a**) Firing rate of a pair of simultaneously recorded deltoid MUs plotted against each other for three different conditions (*columns*) with the deltoid in a lengthened (*blue*) or shorted (*red*) posture. MU157 is slightly more active when the muscle is shortened, while MU158 is much less active. One potential concern is that the change in posture may have altered the spike shape of MU158 such that it is missed in the second (short) posture. Inspection of templates reveals that this was not the case -- both MUs have similar and distinct waveforms across the two postures, as described below. (**b**) *Left*. Template of MU157 across the 5 EMG channels used during this session. *Right*. The 20 waveforms identified in each posture that were most similar to the template. n denotes the total spike counts across all trials in each posture, which increased modestly for MU157 from the long to short posture (even though they decreased for MU158). (**c**) Same as ***b*** for MU158. Taken together, these two panels illustrate that both MUs have distinct waveforms that are not easily confused. In both cases, there was only a slight change in the typical waveform following the move to the short muscle length. In particular, it was not the case that the waveform for MU158 changed and became a poor match with the template.

**Figure S5.**
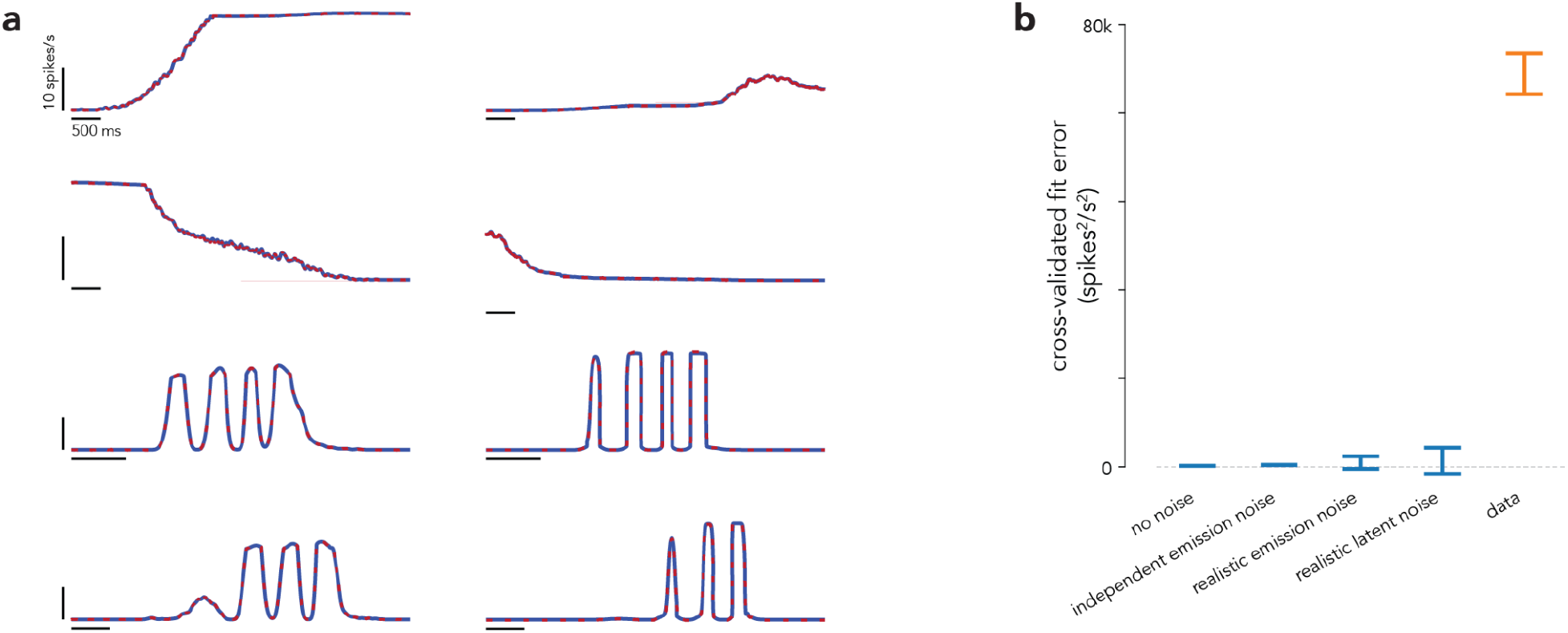
Single-latent model fits to artificial data constructed to be consistent with rigid control, but with various types of noise. We generated realistic simulated data from a 1-latent model by fitting our 1-latent model to the empirical MU response from one session. Then, using the learned latents, link functions, and lags, we generated simulated MU activity. We then fit a different model (with different initialization or parameters) to confirm that it could successfully fit the simulated responses. This acts as both a test of whether optimization succeeds in finding a perfect fit when one is possible, and as a way of documenting the behavior of the cross-validated fit error. **(a)** ‘True’ (*blue traces*) and model fit (*dashed red traces*) responses for two example simulated MUs (*columns*) during four conditions (*rows*). Simulations involved no sampling error. *R*^2^ values of the fit were above 0.99 for all MUs. **(b)** Cross-validated error plots (as in **Fig. 7**) for simulated data for this example session, after incorporating noise into the simulations. ‘Independent emission noise’ is Gaussian noise that is independently added at each time point, with standard deviation for each MU equal to the SEM of MU activity across trials, averaged across time-points and MUs. ‘Realistic emission noise’ is *T*-dimensional gaussian noise (where *T* is the number of time points) generated from a MU’s temporal covariance structure computed across trials (i.e., the covariance matrix of a *T × R* matrix of activity, where *R* is the number of trials). This noise structure is calculated separately for each MU. ‘Realistic latent noise’ is *T*-dimensional gaussian noise that is added to the latent (prior to the link functions), rather than noise directly added to the MU activities. This noise was generated from the latents’ temporal covariance structure computed across trials. Independent noise was added to the latent for each MU, corresponding to each neuron receiving a noisy version of a single latent. Error bars show mean and 95% range of the cross-validation error across multiple partitionings of the data. Note that cross-validated error should be zero on average if sampling noise is the only impediment to a perfect fit, and is thus expected (in that situation) to take on a range of positive and negative values centered near zero. Cross-validated error was indeed near zero for all simulations (and much lower than the fit error for the empirical data, orange) confirming that optimization was successful and cross-validated error behaved as expected. Note that this session had larger single-latent model violations for the empirical data than the average session, so the scale of the y-axis is larger than in **Fig. 7**.

**Figure S6.**
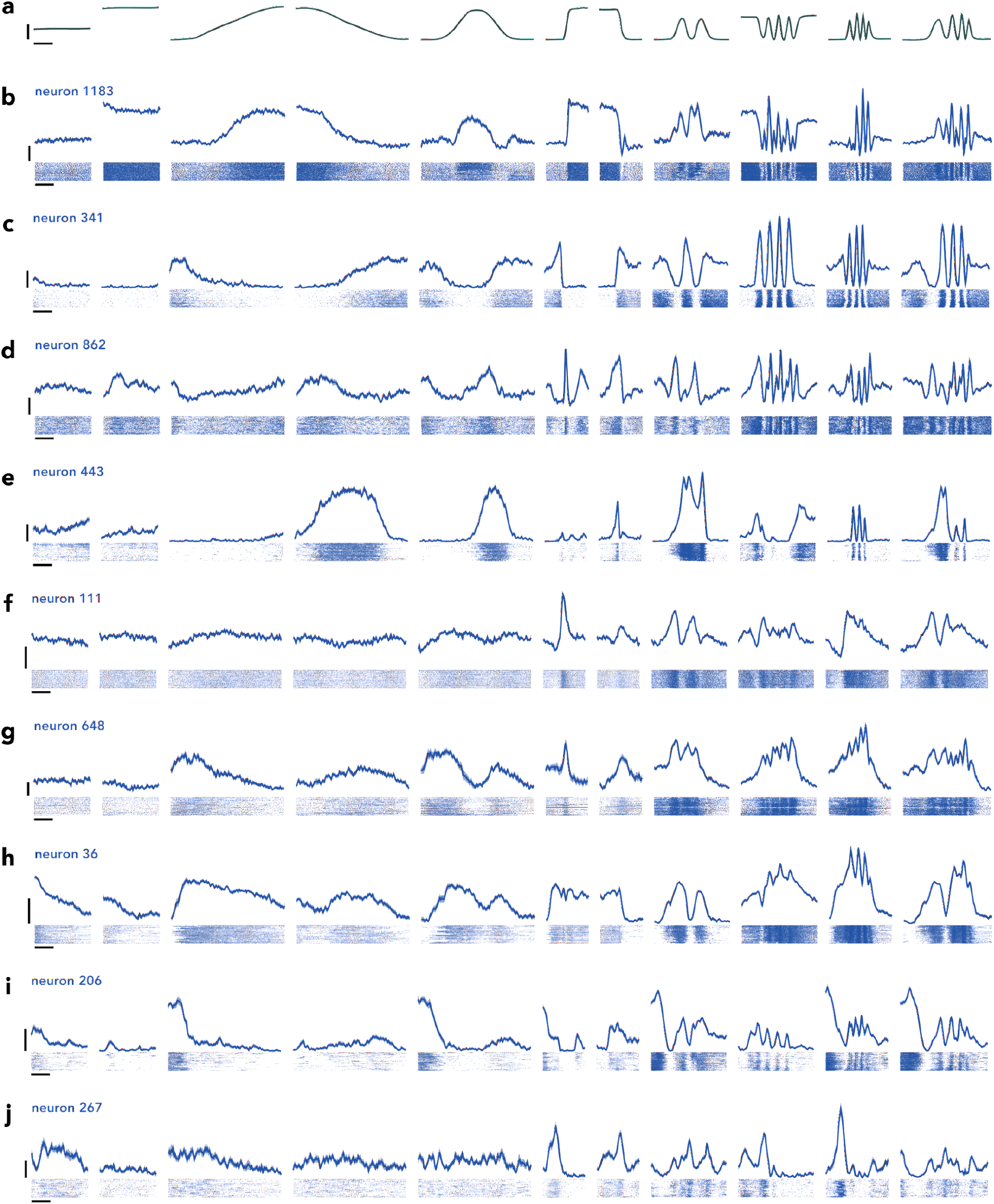
Example M1 responses. (**a**) Trial-averaged forces from one session of dynamic experiments (intermediate static force condition is omitted for space). Vertical scale bar indicates 8 N. Horizontal scale bar indicates 1 s. (**b**-**j**) Trial-averaged firing rates of 9 M1 neurons with standard error (*top*) and single-trial spike rasters (*bottom*). Vertical scale bars indicate 20 spikes/s. Horizontal scale bars indicate 1 s.

## Methods

### Data Acquisition

#### Subject and task

All protocols were in accord with the National Institutes of Health guidelines and approved by the Columbia University Institutional Animal Care and Use Committee. Subject C was an adult, male macaque monkey (*Macaca mulatta*) weighing 13 kg.

During experiments, the monkey sat in a primate chair with his head restrained via surgical implant and his right arm loosely restrained. To perform the task, he grasped a handle with his left hand while resting his forearm on a small platform that supported the handle. Once he had achieved a comfortable position, we applied tape around his hand and velcro around his forearm, which ensured consistent placement within and between sessions. When performing the task, the monkey received both time-varying visual feedback from the screen and time-varying cutaneous feedback from the pressure his hand applied to the handle. There exist reports that adding unusual sensory feedback (e.g., via cutaneous nerve or mechanical stimulation) may alter recruitment under some situations^1^, though this is controversial. This is extremely unlikely to have impacted our results as our task design placed the primary source of feedback (the palm) far from the relevant shoulder muscles, and no interaction across such distance has been suggested (indeed, most studies that report orderly recruitment involve sensory feedback). Furthermore, the handle was held in the same way across force profiles and across stimulation sites, and yet it was often across those conditions that we observed large departures from rigid control.

The handle controlled a manipulandum, custom made from aluminum (80/20 Inc.) and connected to a ball bearing carriage on a guide rail (McMaster-Carr, PN 9184T52). The carriage was fastened to a load cell (FUTEK, PN FSH01673), which was locked in place. The load cell converted one-dimensional (tensile and compressive) forces to a voltage signal. That voltage was amplified (FUTEK, PN FSH03863) and routed to a Performance real-time target machine (Speedgoat) that executed a Simulink model (MathWorks) to run the task. As the load cell was locked in place, forces were applied to the manipulandum via isometric contractions.

The monkey controlled a ‘Pac-Man’ icon, displayed on an LCD monitor (Asus PN PG258Q, 240 Hz refresh, 1920 x 1080 pixels) using Psychophysics Toolbox 3.0. Pac-Man’s horizontal position was fixed on the left hand side of the screen. Vertical position was directly proportional to the force registered by the load cell. For 0 Newtons applied force, Pac-Man was positioned at the bottom of the screen; for the calibrated maximum requested force for the session, Pac-Man was positioned at the top of the screen. Maximum requested forces (see: Experimental Procedures, below) were titrated to be comfortable for the monkey to perform across multiple trials and to activate multiple MUs, but not so many that rendered EMG signals unsortable. On each trial, a series of dots scrolled leftwards on screen at a constant speed (1344 pixels/s). The monkey modulated Pac-Man’s position to intercept the dots, for which he received juice reward. Thus, the shape of the scrolling dot path was the temporal force profile the monkey needed to apply to the handle to obtain reward. We trained the monkey to generate static, step, ramp, and sinusoidal forces over a range of amplitudes and frequencies. We define a ‘condition’ as a particular target force profile (e.g., a 2 Hz sinusoid) that was presented on many ‘trials’, each a repetition of the same profile. Each condition included a ‘lead-in’ and ‘lead-out’ period: a one-second static profile appended to the beginning and end of the target profile, which facilitated trial alignment and averaging (see below). Trials lasted 2.25-6 seconds, depending on the particular force profile. Juice was given throughout the trial so long as Pac-Man successfully intercepted the dots, with a large ‘bonus’ reward given at the end of the trial.

The reward schedule was designed to be encouraging; greater accuracy resulted in more frequent rewards (every few dots) and a larger bonus at the end of the trial. To prevent discouraging failures, we also tolerated small errors in the phase of the response at high frequencies. For example, if the target profile was a 3 Hz sinusoid, it was considered acceptable if the monkey generated a sinusoid of the correct amplitude and frequency but that led the target by 100 ms. To enact this tolerance, the target dots sped up or slowed down to match his phase. The magnitude of this phase correction scaled with the target frequency and was capped at +/- 3 pixels/frame. To discourage inappropriate strategies (e.g., moving randomly, or holding in the middle with the goal if intercepting some dots) a trial was aborted if too many dots were missed (the criterion number was tailored for each condition).

#### Surgical procedures

After task performance stabilized at a high level, we performed a sterile surgery to implant a cylindrical chamber (Crist Instrument Co., 19 mm inner diameter) that provided access to M1. Guided by structural magnetic resonance imaging scans taken prior to surgery, we positioned the chamber surface-normal to the skull, centered over the central sulcus. We covered the skull within the cylinder with a thin layer of dental acrylic. Small (3.5 mm), hand-drilled burr holes through the acrylic provided the entry point for electrodes.

#### Intracortical recordings and microstimulation

Neural activity was recorded with Neuropixels probes. Each probe contained 128 channels (two columns of 64 sites). Probes were lowered into position with a motorized microdrive (Narishige). Recordings were made at depths ranging from 5.6 - 12.1 mm relative to the surface of the dura. Raw neural signals were digitized at 30 kHz and saved with a 128-channel neural signal processor (Blackrock Microsystems, Cerebus).

Intracortical electrical stimulation (20 biphasic pulses, 333 Hz, 400 μs phase durations, 200 μs interphase) was delivered through linear arrays (Plexon Inc., S-Probes) using a neurostimulator (Blackrock Microsystems, Cerestim R96). We did not explore the effectiveness of different parameters but simply used parameters common across many of our studies. We kept pulse-trains relatively short to avoid the more complex movements that occur on longer timescales^2^. Each probe contained 32 electrode sites with 100 μm separation between them. Probes were positioned with a motorized microdrive (Narishige). We estimated the target depth by recording neural activity prior to stimulation sessions. Each stimulation experiment began with an initial mapping, used to select 4-6 electrode sites to be used in the experiments. That mapping allowed us to estimate the muscles activated from each site, and the associated thresholds. Thresholds were determined based on visual observation and were typically low (10-50 μA), but occasionally quite high (100-150+ μA) depending on depth. Across all 32 electrodes, microstimulation induced twitches of proximal and distal muscles of the upper arm, ranging from the deltoid to the forearm. Rarely did an electrode site fail to elicit any response, but many responses involved multiple muscles or gross movements of the shoulder that were difficult to attribute to a specific muscle. Yet some sites produced more localized responses, prominent only within a single muscle head. Sometimes a narrow (few mm^2^) region within the head of one muscle would reliably and visibly pulse following stimulation. Because penetration locations were guided by recordings and stimulation on previous days, such effects often involved the muscles central to performance of the task: the *deltoid* and *triceps*. In such cases, we selected 4-6 sites that produced responses in one of these muscles, and targeted that muscle with EMG recordings. EMG recordings were always targeted to a localized region of one muscle head (see below). In cases where stimulation appeared to activate only part of one muscle head, EMG recordings targeted that localized region.

#### EMG recordings

Intramuscular EMG activity was recorded acutely using paired hook-wire electrodes (Natus Neurology, PN 019-475400). Electrodes were inserted ~1 cm into the muscle belly using 30 mm x 27 G needles. Needles were promptly removed and only the wires remained in the muscle during recording. Wires were thin (50 um diameter) and flexible and their presence in the muscle is typically not felt after insertion, allowing the task to be performed normally. Wires were removed at the end of the session.

We employed several modifications to facilitate isolation of MU spikes. As originally manufactured, two wires protruded 2 mm and 5 mm from the end of each needle (thus ending 3 mm apart) with each wire insulated up to a 2 mm exposed end. We found that spike sorting benefited from including 4 wires per needle (i.e., combining two pairs in a single needle), with each pair having a differently modified geometry. Modifying each pair differently meant that they tended to be optimized for recording different MUs^3^; one MU might be more prominent on one pair and the other on another pair. Electrodes were thus modified as follows. The stripped ends of one pair were trimmed to 1 mm, with 1 mm of one wire and 8 mm of the second wire protruding from the needle’s end. The stripped ends of the second pair were trimmed to 0.5 mm, with 3.25 mm of one wire and 5.25 mm of the second wire protruding. Electrodes were hand fabricated using a microscope (Zeiss), digital calipers, precision tweezers and knives. During experiments, EMG signals were recorded differentially from each pair of wires with the same length of stripped insulation; each insertion thus provided two active recording channels. Four insertions (closely spaced so that MUs were often recorded across many pairs) were employed, yielding eight total pairs. The above approach was used for both the dynamic and muscle-length experiments, where a challenge was that normal behavior was driven by many MUs, resulting in spikes that could overlap in time. This was less of a concern during the microstimulation experiments. Stimulation-induced responses were typically fairly sparse near threshold (a central finding of our study is that cortical stimulation can induce quite selective MU recruitment). Thus, microstimulation experiments employed one electrode pair per insertion and 8 total insertions (rather than two pairs and 4 total insertions), with minimal modification (exposed ends shorted to 1 mm).

Raw voltages were amplified and analog filtered (band-pass 10 Hz - 10 kHz) with ISO-DAM 8A modules (World Precision Instruments), then digitized at 30 kHz with a neural signal processor (Blackrock Microsystems, Cerebus). EMG signals were digitally band-pass filtered online (50 Hz - 5 kHz) and saved.

#### Experimental procedures

Cortical recordings were performed exclusively during one set of experiments (‘dynamic’, defined below), whereas EMG recordings were conducted across three sets of experiments (dynamic, ‘muscle length’, and microstimulation). In a given session, the eight EMG electrode pairs were inserted within a small (typically ~2 cm^2^) region of a single muscle head. This focus aided sorting by ensuring that a given MU spike typically appeared, with different waveforms, on multiple channels. This focus also ensured that any response heterogeneity was due to differential recruitment among neighboring MUs.

In dynamic experiments, the monkey generated a diverse set of target force profiles. The manipulandum was positioned so that the angle of shoulder flexion was 25° and the angle of elbow flexion was 90°. Maximal requested force was 16 Newtons. We employed twelve conditions (**Supp Fig. 1**) presented interleaved in pseudo-random order: a random order was chosen, all conditions were performed, then a new random order was chosen. Three conditions employed static target forces: 33%, 66% and 100% of maximal force. Four conditions employed ramps: increasing or decreasing across the full force range, either fast (lasting 250 ms) or slow (lasting 4 s). Four conditions involved sinusoids at 0.25, 1, 2, and 3 Hz. The final condition was a 0-3 Hz chirp. The amplitude of all sinusoidal and chirp forces was 75% of maximal force, except for the 0.25 Hz sinusoid, which was 100% of maximal force. Recordings in dynamic experiments were made from the deltoid (typically the anterior head and some from the lateral head) and the triceps (lateral head).

In muscle-length experiments, the monkey generated force profiles with his deltoid at a long or short length (relative to the neutral position used in the dynamic experiments). The manipulandum was positioned so that the angle of shoulder flexion was 15° (long) or 50° (short), while maintaining an angle of elbow flexion of 90°. Maximal requested forces were 18 N (long) and 14 N (short). Different maximal forces were employed as it appeared more effortful to generate forces in the shortened position. To ensure enough trials per condition, we employed only a subset of the force profiles used in the dynamics experiments. These were 2 static forces (50% and 100% of maximal force), the slow increasing ramp, both increasing and decreasing fast ramps, all four sinusoids and the chirp. These were presented interleaved in pseudorandom order for multiple trials (~30 per condition) for the lengthened position (15°) before changing to the shortened position (50°). In most experiments we were able to revert to the lengthened position (15°) at the end of the session, and verify that MU recruitment returned to the originally observed pattern. Recordings in muscle-length experiments were made from the *deltoid* (anterior head).

Microstimulation experiments employed recordings from the lateral deltoid and lateral triceps. Both these muscles exhibited strong task-modulated activity, as documented in the dynamic and muscle-length experiments. We also included recordings from the sternal pectoralis major, which showed only modest task-modulated activity, as we found cortical sites that reliably activated it. The manipulandum was positioned so that the angle of shoulder flexion was 25° and the angle of elbow flexion was 90° (as in dynamic experiments). Maximal force was typically set to 16 N, but was increased to 24 N and 28 N for two sessions each in an effort to evoke greater muscle activation. Microstimulation experiments employed a limited set of force profiles: four static forces (0, 25%, 50% and 100%), and the slow (4 s) increasing ramp. The ramp was included to document the natural recruitment pattern during slowly changing forces. Microstimulation was delivered once per trial during the static forces, at a randomized time (1000-1500 ms relative to when the first dot reached Pac-Man). Because stimulation evoked activity in muscles used to perform the task, it sometimes caused small but detectable changes in force applied to the handle. However, these were so small that they did not impact the monkey’s ability to perform the task and appeared to go largely unnoticed. These experiments involved a total of 17-25 conditions: the ramp condition (with no stimulation) plus the four static forces for the 4-6 chosen electrode sites. These were presented interleaved in pseudorandom order.

### Data Processing

#### Signal processing and spike sorting

Cortical voltage signals were spike sorted using KiloSort 2.0^4^ A total of 881 neurons were isolated across 15 sessions.

EMG signals were digitally filtered offline using a second-order 500 Hz high-pass Butterworth. Any low SNR or dead EMG channels were omitted from analyses. Motor unit (MU) spike times were extracted using a custom semi-automated algorithm. As with standard spike-sorting algorithms used for neural data, individual MU spikes were identified based on their match to a template: a canonical time-varying voltage across all simultaneously recorded channels (example templates are shown in **Fig. S1**, *bottom left*). Spike templates were inferred using all the data across a given session. A distinctive feature of intramuscular records (compared to neural recordings) is that they have very high signal-to-noise (peak-to-peak voltages on the order of mV, rather than uV, and there is negligible thermal noise) but it is common for more than one MU to spike simultaneously, yielding a superposition of waveforms. This is relatively rare at low forces but can become common as forces increase. Our algorithm was thus tailored to detect not only voltages that corresponded to single MU spikes, but also those that resulted from the superposition of multiple spikes. Detection of superposition was greatly aided by the multi-channel recordings; different units were prominent on different channels. Further details are provided in the Supplementary Methods. We used constructed data with realistic properties (e.g., recruitment based on the actual force profiles we used, and waveforms based on actual recorded waveforms) to verify that spike-sorting was accurate in circumstances where the ground truth was known. Mis-sorted spikes were rare, and they were similarly rare during slowly changing forces versus high-frequency sinusoids. Spike-sorting errors are thus unlikely to explain differences in the empirical recruitment between these situations.

#### Trial alignment and averaging

Single-trial spike rasters, for a given neuron or MU, were converted into a firing rate via convolution with a 25 ms Gaussian kernel. One analysis (**Fig. 4d**) focused on single-trial responses, but most employed trial-averaging to identify a reliable average firing rate. To do so, trials for a given condition were aligned temporally and the average firing rate, at each time, was computed across trials. Stimulation trials were simply aligned to stimulation onset. For all other conditions, each trial was aligned on the moment the target force profile ‘began’ (when the target force profile, specified by the dots, reached Pac-Man). This alignment brought the actual (generated) force profile closely into register across trials. However, because the actual force profile could sometimes slightly lead or lag the target force profile, some modest across-trial variability remained. Thus, for all trials with changing forces, we realigned each trial (by shifting it slightly in time) to minimize the mean squared error between the actual force and the target force profile. This ensured that trials were well-aligned in terms of the actual generated forces (the most relevant quantity for analyses of MU activity). Trials were excluded from analysis if they could not be well aligned despite searching over shifts from −200 to 200 ms.

### Data Analysis

#### Interaction effects of microstimulation

We quantified whether microstimulation-evoked responses displayed an interaction effect between stimulation electrode and MU identity using a two-way ANOVA. For each session and static force level, we computed the largest absolute change between the mean firing rate during a baseline period and the firing rate at each time during a response period for each MU and stimulation trial. Baseline and response periods were defined as −300-0 ms and 30-130 ms with respect to stimulation onset. We performed a two-way ANOVA (anovan, MATLAB) using the vector of single-trial responses and a set of two grouping vectors containing the identity of each MU and stimulation electrode. To evaluate whether the responses for a particular session displayed an interaction effect, p-values were divided by the number of static force amplitudes.

#### Quantifying motor unit flexibility

We developed two analyses that quantified MU-recruitment flexibility without directly fitting a model (model-based quantification is described below). These two analyses were used to produce the results in **Figures S2** and **S3**, respectively. Both methods leverage the definition of rigid control to detect population-level patterns of activity that are inconsistent with rigid control even under the most generous of assumptions.

Let **r***_t_* = [*r*_1,*t*_ *r*_2,*t*_… *r_n,t_*]^⊤^ denote the population state at time *t*, where *r_i,t_* denotes the firing rate of the *i*^th^ MU. If **r***_t_* traverses a 1-D monotonic manifold, then as the firing rate of one MU increases, the firing rate of all others should either increase or remain the same. More generally, the change in firing rates from *t* to *t′* should have the same sign (or be zero) for all MUs. If changes in firing rate are all nonnegative (increasing or staying the same), then one can infer that common drive increased from *t* to *t′*. If the changes in firing rate are all nonpositive (decreasing or staying the same), then one can infer that common drive decreased. Both these cases (all nonnegative or all nonpositive) are consistent with rigid control because there exists some 1-D monotonic manifold that contains the data at both *t′* and *t*.

On the other hand, departures from a 1-D monotonic manifold can be inferred as moments when the firing rates of one or more MUs increase as others’ decrease. Both our analyses seek to quantify the magnitude of such departures while being very conservative. Specifically, the size of a departure was always measured as the smallest possible discrepancy from a 1-D manifold, based on all possible 1-D manifolds. To illustrate the importance of this conservative approach, consider a situation where the firing rate of MU1 increases considerably while MU2’s rate decreases slightly from *t* to *t′*. This scenario would be inconsistent with activity being modulated solely by a common input, yet it would be impossible to know which MU reflected an additional or separate input. Perhaps common drive decreased slightly (explaining the slight decrease in MU2’s rate) but MU1 received an additional large, private excitatory/inhibitory input. This would indicate a large departure from rigid control. Yet another possibility is that common drive increased considerably (explaining the large increase in MU1’s rate) and that MU2’s rate failed to rise because it was already near maximal firing rate. This would not explain why MU2’s rate went down, but if that decrease was small it could conceivably be due to a very modest departure from idealized rigid control. Thus, to be conservative, one should quantify this situation as only a slight deviation from the predictions of rigid control. Both methods described below were designed to do so; when MU activities were anticorrelated, we identified the largest increase and decrease in firing rates, then reported the change that was smaller in magnitude.

For the first analysis, we computed the largest nonnegative change in firing rates from *t* to *t′* for a population of *n* MUs as

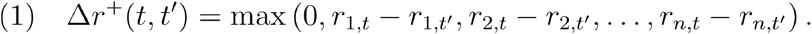

If a 1-D monotonic manifold can be drawn through **r***_t_* and **r***_t′_*, then either Δ*r*^+^(*t, t′*) or Δ*r*^+^(*t′, t*) will be zero. Otherwise, Δ*r*^+^(*t, t′*) will capture the largest increase (across MUs) in rate from *t* to *t′* while Δ*r*^+^(*t′, t*) will capture the largest decrease.Thus, we computed departures from a monotonic manifold at the level of an individual MU as

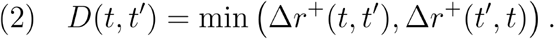

As examples, consider a population of two MUs with **r***_t_* = [10, 10] and **r***_t′_* = [15, 25], These states would be consistent with an increase in common drive from *t* to *t′*, so *D*(*t, t′*) = 0 (**Supp Fig. S2a**, *left*). Conversely, **r***_t_* = [10, 10] and **r***_t′_* = [9, 30] (*center*) suggests a violation of rigid control, but that violation might be small; one can draw a manifold that passes through [10, 10] and comes within 1 spike/s of [9, 30]. In this case, *D*(*t, t′*) = 1. Finally, **r***_t_* = [10, 10] and **r***_t′_* = [0, 30] (*right*) argue for a sizable violation; [0, 30] is at least 10 spikes/s distant from any monotonic manifold passing through [10, 10], so *D*(*t, t′*) = 10.

It is worth emphasizing that (**eq. 2**) can readily be computed for a population with more than two MUs, but the analysis ultimately reduces to a comparison of two MUs: one whose firing rate increased the most and the other whose firing rate decreased the most across a pair of time points.

To extend our analysis to multiple time points, we computed the ‘MU displacement’ as

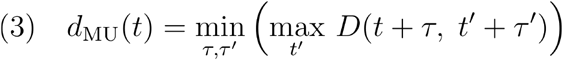

where *t′* indexes over all other times and conditions, and *τ* and *τ′* are time lags. The inclusion of time lags ensures that departures from a monotonic manifold cannot simply be attributed to modest differences in response latencies across MUs. In our analyses, we optimized over *τ τ′* ∈ [−25, 25] ms. *d*_MU_ is exceedingly conservative; it makes no assumptions regarding the manifold other than that it is monotonic, and identifies only those violations that are apparent when comparing just two times. When analyzing a single session we considered the distribution of *d*_MU_(*t*) across all times *t*. When considering all sessions, we simply took the maximum within each session.

An advantage of the *d*_MU_ metric is interpretational simplicity; it identifies pairs of times where the joint activity of two MUs cannot lie on a single 1-D monotonic manifold. A disadvantage is that it does not also capture the degree to which multiple other MUs might also have activity inconsistent with a 1-D monotonic manifold. To do so, we employed a second metric that quantifies MU-recruitment flexibility at the population level. Under the assumptions of rigid control, the magnitude of common drive determines the population state and therefore the summed activity of all MUs or, equivalently, its L1-norm, ║r║_1_. Increases and decreases in common drive correspond, in a one-to-one manner, to increases and decreases in ║r║_1_. Violations of rigid control can thus be inferred if a particular norm value, λ, is associated with different population states. Geometrically, this corresponds to the population activity manifold intersecting the hyperplane defined by ║r║_1_ = λ at multiple locations.

We thus defined the motor neuron pool (MNP) dispersion as

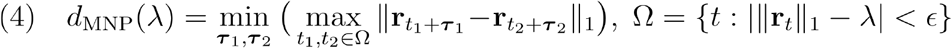

where ***τ***_1_, ***τ***_2_ are time lag vectors, of the same dimensionality as **r**, and *ϵ* is a small constant. Conceptually, the dispersion identifies the pair of time points when the population states are the most dissimilar, while having norms within *ϵ* of λ. As when computing *d*_MU_(*t*) we minimized *d*_MNP_(λ) over time lags so as to only consider dispersions that could not be simply attributed to latency differences across MUs. For our analyses, we set *ϵ* = 1 and optimized over ***τ***_1_, ***τ***_2_ ∈ [−25, 25] ms. When analyzing a single session we considered*d*_MNP_(λ) as a function of λ. When considering all sessions, we simply took the maximum within each session.

We compared displacement and dispersion values for the true MU population and an artificial population (**Figs. 3c**, **5j, k**, **6h, i**). The artificial population was generated so as to resemble the empirical responses as closely as possible, while being describable by a single latent factor plus corruption by sampling error (i.e., it obeys rigid control except for the addition of realistic sampling error). To make the artificial population response consistent with rigid control, we extracted the first principal component (PC1) from the trial-averaged firing rates over all conditions (any dimension would be reasonable, but PC1 is a sensible choice as it presumably would capture the most variance for data that did obey rigid control). For each individual trial, we projected the population firing rate (for all times during that trial) onto PC1, reconstructed the firing rate of each MU, then rectified to remove any negative rates. This resulted in data that perfectly obeyed rigid control (and would have resulted in both displacement and dispersion values of exactly zero). To reintroduce sampling error, we redrew trials (as many as were originally recorded) in which the population response on a given redrawn trial was composed of individual MU responses on different (randomly and independently drawn) trials. For example, the population response on the first redrawn trial might contain the response of MU1 on trial 8, the response of MU2 on trial 33, etc. Thus, each redrawn trial was a noisy (with noise matching the original recordings) observation of an underlying population response that obeyed rigid control. Trial averages were then computed to generate the artificial population response. We then measured displacement and dispersion. This process was repeated ten times with different random draws. There are thus ten times as many red traces (in Fig. 3c, Fig 5j,k, and Fig. 6h,i) as sessions.

Computation of displacement and dispersion requires that one choose a set of population responses spanning both time and one or more conditions. The values of *t* and *t′* index that set. Depending on the analysis, *t* and *t′* may be constrained to lie within the same condition, or may be allowed to span both conditions. For example, in **Fig. 5m**, the purple trace plots the distribution of values of *d*_MU_(*t*) when both *t* and *t′* are constrained to lie within the set of population responses observed during the slow four-second ramp. The green trace plots the distribution of values when *t* and *t′* can span the full set of population responses observed during all force profiles. Many analyses use this strategy of computing displacement and dispersion for both a more restrictive set of conditions and a more expansive set. For example, Fig. 3b computes displacement both when considering each stimulation site on its own (1-stim) and when considering the population response for all (all-stim). For 1-stim, the maximum displacement is computed separately for each stimulation site, with *t* and *t′* indexing within the response for that site. The maximum is then taken across all values of *t*. Because there are multiple sites, there are multiple such maxima and we report the largest. For all-stim, *t* and *t′* index across responses for all stimulation sites, and the maximum is taken across *t*.

#### Optimal MU recruitment model

We developed a computational model to predict the optimal recruitment strategy for generating a particular force profile by an idealized motor pool. MU twitch responses were simulated using a standard model, in which peak tension and contraction time are inversely related (i.e., MUs with smaller peak tensions have longer contraction times)^5^. The set of MU firing rates for generating a target force profile was derived as the solution that minimized a cost function that depended principally on the mean-squared error between the target and simulated forces (in addition to modest regularizers). Minimizing this cost function is a convex optimization problem, which we solved numerically. For details, see Supplementary Materials.

#### Latent factor model formulation

We developed a probabilistic latent variable model of MU activity. Let *y_i,t_* be the known activity of the *i*^th^ MU at time *t*. Let *x_t_* be the unknown latent variables at time *t*, which are shared between all MUs. We can fit this model with one latent (Fig. 7; *x_t_* can be a single value) or multiple latents (Fig. 8). The relationship between the MU activity and the underlying latent(s) is given by:

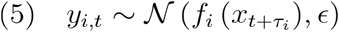

where *f_i_* denotes a learned link function for the *i*^th^ MU and *τ_i_* denotes a learned lag between its response and the shared latent variables. We constrained *τ_i_* ∈ [−25, 25] ms. Note that the absolute values of these are meaningless (because the timing of the inferred latent is unconstrained) -- what is important is the range, which we set so that the largest allowable difference in lag between two MUs is 50ms. 50 ms is larger than the likely range of latencies due to variable conduction velocities, but not so large as to be wildly unrealistic. This accords with our general strategy of allowing the model as much reasonable expressivity as possible so that interpretation is conservative.

To allow expressive, monotonically increasing link functions with nonnegative outputs, we parameterized *f_i_* as a rectified monotonic neural network. We fit each *f_i_* using a two-hidden-layer feedforward neural network, in which the weights were constrained to be positive. This class of network with only a single hidden layer is able to model any univariate monotonic function (as is the case when our model uses a single latent variable)^6^. The positivity constraint was achieved by letting each weight *w* = ln(1 + *e^u^*), where the values of *u* were fit within the model. For the models with one latent variable (**Fig. 7**), we employed 10 hidden units per layer. This allows considerably more expressivity than the commonly used sigmoid, while still encouraging relatively smooth monotonic functions. Performance improved negligibly if we increased the number of hidden units up to 100 per layer, allowing quite non-smooth functions. Thus, the inability of the single-latent model to account for the data was not due to insufficiently expressive link functions. (This inference can also be made by inspecting the data, which often depart from a monotonic manifold and thus cannot be fit by a single-latent model with any monotonic link functions.) Early versions of our model used logistic or spline-based link functions and led to the same scientific result: a single-latent model cannot sufficiently explain MU activity across conditions. We used the neural-network-based link functions simply to be conservative and confirm that results held when expressive link functions were allowed. During model training, the output of the neural network was passed through a ‘leaky rectified linear unit’ with a slope value of 0.01 for negative inputs to the leaky ReLU. After training was completed, we used standard rectification on the output.

When fitting models with multiple latent variables (Fig. 8), we used 20 hidden units per layer, as this improved performance of multi-latent models. Link functions were thus highly expressive. We therefore stress that the analysis likely underestimates the number of latents necessary to fit the data (i.e., more latents would likely be needed with less expressive link functions). This tradeoff highlights a limitation of the single-latent model. It is ideally suited to testing the hypothesis of rigid control. However, once that hypothesis is rejected, meaningfully extending the model requires incorporating constraints based on physiology or anatomy, and many of those constraints are presently unknown.

Optimization sought to minimize the mean squared error (MSE) between model fits and empirical population firing rates. Given that rates were derived from spikes, another option would have been to maximize spiking likelihood under a Poisson model. We chose to minimize MSE for three reasons. First, MU spiking has very non-Poisson statistics. Second, we wished the model to be able to fit both single-trial rates and trial-averaged rates, and MSE is very natural in the latter case as sampling error should become fairly Gaussian due to the central limit theorem. Third, in our central analysis (Fig. 7c), squared error provides an intuitive metric of the distance between what the data does and what the model can explain. Furthermore, squared error can be computed in a manner that is not biased upward by sampling error that inevitably arises due to spiking variability (by cross-validating across partitions, see below). A core element of our strategy was always to give the model every chance to succeed (only then is it fair to reject it following failure). Thus, it was most appropriate to minimize the metric by which success would be judged. We confirmed that model residuals were close to Gaussian distributed in circumstances where the model was expected to fit well (e.g., during slowly changing forces).

For our example model fits (Fig. 7b), we wished to plot trajectories with smoothness that were reasonably realistic (i.e., on the same scale as the data), without hurting the ability of the model to fit. We thus incorporated additional smoothness by letting 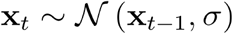, where smaller values of *σ* encouraged greater temporal smoothness. We set *σ* to 0.01 for this figure. Greater temporal smoothness leads to more realistic trajectories, which are more easily compared with the empirical trajectories (which had of course been minimally smoothed). However, quantitatively, results were nearly identical regardless of whether we did or did not require smoothness in the latent variable. Thus, for simplicity, our quantitative analyses did not insist on any additional smoothness. This corresponds to allowing the latent variable (i.e., the common drive) to change very swiftly if that helped fit the data. This choice is conservative because, if the model still fails to fit, it cannot be because we imposed smoothness.

The standard conception of rigid control is that a single latent variable (a common-drive force command) determines MU activity via MU-specific link functions (providing each MU with its own threshold etc.). While those link functions are typically modeled as static, some MUs exhibit history dependence due to persistent inward currents (PICs) that increase activity. PICs rarely alter recruitment order because they are most prevalent in small MUs, which for a given force level are the more active MUs to begin with (making them even more active does not alter recruitment order). However, because PICs cannot be captured well by a fixed link function, it is possible that they are a partial source of model failure. They are unlikely to be a major source error for two reasons. First, inspection reveals many model failures that are unrelated to PICs (e.g., **Fig. 7b**). Second, we saw putative PIC-based effects only occasionally. Still, the fact that they were sometimes observed means that it is likely that model fits suffered modestly because they could not fit PIC-based effects. To control for this, we fit models in which, rather than giving units fixed lags, we allowed the mapping between the latent and the MU activity to have internal dynamics (accomplished by adding a hidden recurrent unit, unique for each MU, between the latent and the MU’s feedforward neural network link function). Despite the increase in expressivity the same results were observed: the model fit well when fitting slowly changing forces but fit poorly when fitting all conditions together. In the latter case, total error was reduced by ~25% as a result of allowing dynamics, but was still high. Inspection revealed that some of the reduction in error was indeed due to the model now being able to account for effects that were likely PIC-driven. However, these were not the principal source of model error.

#### Latent factor model fitting

To infer the most likely distribution of latent variables given the data (i.e., the model posterior, *p*(x|y)), and to learn the link functions, we used variational inference with a mean-field approximation for the posterior approximation. As an inference method, we used black-box variational inference^7,8^, which performs gradient-based optimization via automatic differentiation to maximize the model’s evidence lower bound, in order to directly update both the mean and variance of the variational posterior, and the model parameters. We used the Adam algorithm for optimization^9^. We iterated between (1) optimizing the posterior and link functions while holding response lags fixed, as described above and (2) optimizing the response lags. As the response lags were integers rather than continuous variables, rather than using a gradient-based method, during each iteration of updating the lags, we selected lags, within 2 ms of the previous iteration’s lags, that maximized the data’s likelihood. Using simulated data, we confirmed that this approach accurately inferred known lags. Post model-fitting, when predicting MU activity, we used the mean of the posterior distribution as the latent input at each time.

To initialize the model fits, we set the mean of the variational distribution to the mean activity (across neurons) at each time. We then initialized the link functions by fitting the monotonic neural network to predict MU firing rates from this mean activity across time. Lags were initialized at 0 ms for all MUs. This initialization procedure led to very stable model predictions across multiple model fits, and thus all analyses were done using single model fits (as opposed to choosing the best among many model fits).

Prior to fitting the model, the firing rate of each MU was normalized by its maximum response across conditions. Normalization did not alter the ability of the model to fit the data, but simply encouraged the model to fit all MUs, rather than just the high-rate units. Additionally, the likelihood of each time point was weighted by the duration of the experimental condition, so that each condition mattered equally within the model regardless of duration. When fitting to single trials, we also weighted each condition by its trial count, again so that each condition had equal importance. All model fits were done within individual sessions.

#### Quantifying residual model errors

To compute the cross-validated model residuals, we first randomly split the single-trial firing rates for each MU into halves, and computed the trial-average responses for each half: ***y***_*i*,1_ and ***y***_*i*,2_. We then fit the latent variable model to each half, which yielded a pair of predicted responses, 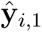 and 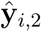 The cross-validated model residuals were calculated as the dot product between the residual errors of each half: 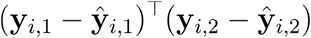. We computed the median cross-validated residuals across all MUs and sessions for a given partitioning of the data. The above steps were then repeated for 10 different random splits of trials and we reported the mean +/- standard error of the median error across re-partitions and fits.

As a control (**Fig. 7c**), we modified the data so that a single latent variable could fully account for all responses. To do so, we reconstructed the firing rates using only the first principle component of the trial average firing rates. For example, if w is the *n* × 1 loading vector for the first principal component, then *Y*_1_ the *ct × n* matrix of responses for one partitioning of the data, was reconstructed as [*Y*_1_**ww**^⊤^]_+_, where the rectification ensures that all firing rates are non-negative. Using these reconstructed firing rates, we performed the same residual error analysis. Because of the rectification, the modified data are not one-dimensional in the linear sense (there would be multiple principal components with non-zero variance). Yet because the data will lie on a one-dimensional monotonic manifold, cross-validated error should be near zero when fitting the model, which is indeed what we observed. This confirms that optimization succeeds in finding an essentially perfect fit when the data lie within the scope of things that can be accounted for by the model. We repeated this control, with the same result, with artificial data generated in other ways and with different amounts of sampling error (**Fig. S5**).

#### Consistency plots

We fit the model to the activity of single trials. We aimed to determine whether, when fit to two conditions, the model consistently overestimated the true firing rates in one condition and underestimated the firing rates in the other condition. To do so, we calculated the mean model error across time on every trial for each condition. Let *E*(1, *tr*) and *E*(2, *tr*) denote the mean errors for a particular MU, pair of conditions (indexed by 1 and 2), and trial *tr*. We calculated the consistency for the MU and conditions as

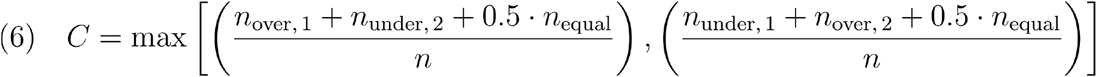

where

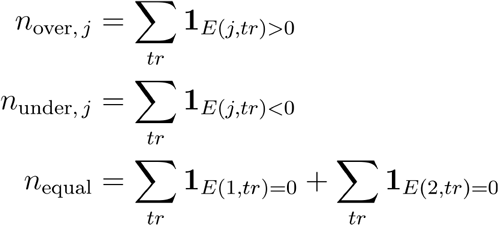

*n* is the total number of trials across both conditions, and **1***_A_* is the indicator function (1 if *A* is true; 0 otherwise). **Eq. (6)** determines the fraction of times one condition had negative errors and the other had positive errors, while accounting for trials with no error. Prior to performing this consistency calculation, we set all *E*(*j, tr*) with absolute value less than 0.01 to 0, so that the sign of negligible errors was not considered. We also removed *E*(1, *tr*) or *E*(2, *tr*) in which the MU had zero actual and predicted activity, because it was impossible for the predicted activity to undershoot the true activity in this setting.

We calculated the fraction of MUs that had *C* > 0.8 and an average error of at least 0.01 across trials (to ensure that outlier trials did not lead to false positives of consistent errors). We excluded MUs who had zero activity in > 80% of trials in the two conditions being analyzed. Consequently, the number of MUs included in the analysis) varied for each pair of conditions.

To calculate a chance-level baseline, for each MU, we calculated the probability that greater than 80% of the included trials would have a positive or negative error, assuming that each trial has an independent 50/50 chance of being positive or negative. More precisely, let *F*(*k; n, p*) be the cumulative density function of a binomial distribution of having *k* successes in *n* Bernoulli events, each event with probability *p* of being a success. We calculate *p_i_* = 2(1 – *F*(ceil[0.8*_n_i__*]; *n_i_*, 0.5)), where *n_i_* is the number of total trials included for MU *i* and 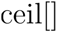 gets the next integer. The total expected fraction of MUs with *C* > 0.8 by chance is thus 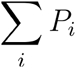.

#### Cross-validated reliability dimensionality estimate

To estimate the dimensionality of M1, we randomly split the single-trial firing rates for each neuron into two groups and averaged over trials within each group. Let *Y*_1_ and *Y*_2_ denote the *CT × N* matrices of trial-averaged responses for each partition (*CT* condition-times and *N* neurons). Let **w***_i_* (an *N* × 1 vector) denote the *i*^th^ principal component (PC) of *Y_i_*. The reliability of PC *i* was computed as the correlation between *Y*_1_**w***_i_* and *Y*_2_**w***_i_*. We repeated this process for 25 re-partitions over trials to obtain confidence intervals. Our method is inspired by Churchland et al.^10^ and conceptually similar to but distinct from the cross-validated PCA analysis of Stringer *et al*., which estimates the stimulus-related (‘signal’) neural variance based on spontaneous activity across many neurons on single trials^11^. PCA was used simply because it concentrates variance in as few dimensions as possible, which is conservative given the present goals.

To create simulated data sets with dimensionality *k*, we computed 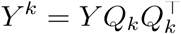, where *Y* is the matrix of M1 firing rates averaged over all trials, and *Q_k_* denotes the first *k* columns of a random orthonormal matrix. Simulated single-trial spikes were generated for each neuron using an inhomogeneous Poisson process with rate given by the corresponding column of *Y_k_*. Simulated spikes were smoothed using a 25 ms Gaussian kernel, and the cross-validated reliability metric applied as described above.

